# Interrogation of the Dynamic Properties of Higher-Order Heterochromatin Using CRISPR/dCas9

**DOI:** 10.1101/2021.06.06.447300

**Authors:** Yuchen Gao, Mengting Han, Stephen Shang, Haifeng Wang, Lei S. Qi

## Abstract

Eukaryotic chromosomes feature large regions of compact, repressed heterochromatin hallmarked by Heterochromatin Protein 1 (HP1). HP1 proteins play multi-faceted roles in shaping heterochromatin, and in cells, HP1 tethering to individual gene promoters leads to epigenetic modifications and silencing. However, emergent properties of HP1 at supranucleosomal scales remain difficult to study in cells due to lack of appropriate tools. Here, we develop CRISPR-Engineered Chromatin Organization (EChO), combining live cell CRISPR imaging with inducible large-scale recruitment of chromatin proteins to native genomic targets. We demonstrate that human HP1α tiling across kilobase-scale genomic DNA forms novel contacts with natural heterochromatin, integrates two distantly targeted regions, and reversibly changes chromatin from a diffuse to compact state. The compact state exhibits delayed disassembly kinetics and represses transcription across over 600 kilobases. These findings support a polymer model of HP1α-mediated chromatin regulation and highlight the utility of CRISPR-EChO in studying supranucleosomal chromatin organization in living cells.

## INTRODUCTION

The separation of the genome into compact, transcriptionally repressed heterochromatin and diffuse, transcriptionally active euchromatin represents a fundamental mechanism of chromatin organization in eukaryotes. On linear chromosomes, heterochromatin is predominantly found at repetitive, gene poor sequences near the centromeres and telomeres, while euchromatin is present at unique, gene rich chromosome arms (Wallrath et al., 2014). Within the 3-D nucleus, the two forms of chromatin are observed by microscopy to occupy spatially distinct regions (Politz et al., 2013), and are observed in high throughput chromosome conformation capture (Hi-C) studies to preferentially self-associate into mutually exclusive compartments (Lieberman-Aiden et al., 2009; Rao et al., 2014). A central question in chromatin biology is to understand the factors at different regulatory scales that determine the spatial, structural, and functional organization of euchromatin and heterochromatin.

Heterochromatin Protein 1 (HP1) is a major component of heterochromatin and possesses a multi-faceted role in shaping the properties of heterochromatin. HP1 is a protein family that is highly conserved across eukaryotes from yeast to humans, and in humans consists of the orthologs HP1α, HP1β, and HP1γ (Lomberk et al., 2006). *In vitro*, HP1 orthologs can form oligomerized assemblies (Canzio et al., 2011; Larson et al., 2017; Yamada et al., 1999), bridge and compact nucleosomes (Azzaz et al., 2014; Machida et al., 2018), and promote liquid-liquid phase separation (Larson et al., 2017; Sanulli et al., 2019; Strom et al., 2017). These properties suggest that HP1 actively contributes to supranucleosomal organization of heterochromatin but have been challenging to study in cells. *In vivo*, the causal role of HP1 for regulating chromatin has been investigated by using synthetic DNA binding domains to tether HP1 family members at targeted synthetic or endogenous euchromatic sites (Braun et al., 2017; Danzer and Wallrath, 2004; Hathaway et al., 2012; Li et al., 2003; Seum et al., 2001; Verschure et al., 2005). These studies found that HP1 tethering can induce local epigenetic remodeling such as H3K9 methylation (Braun et al., 2017; Hathaway et al., 2012; Verschure et al., 2005), DNA methylation (Hathaway et al., 2012), and H3K4 demethylation (Hathaway et al., 2012) and, when tethered at gene promoters, can induce silencing of the adjacent gene (Braun et al., 2017; Danzer and Wallrath, 2004; Hathaway et al., 2012; Li et al., 2003). However, nearly all existing HP1 tethering studies target HP1 to a small region within a single gene regulatory element, while natural heterochromatin exists along megabase-scale regions of the genome and occupies large parts of the 3-D nucleus. Therefore, while the function of individual HP1 proteins may be captured, any emergent properties of HP1 acting collectively at higher scales to regulate chromatin structure and behavior are missed.

CRISPR/Cas9 technologies have revolutionized the way in which chromatin organization can be studied at varying scales. At individual gene regulatory elements, nuclease-deactivated Cas9 (dCas9)-based transcriptional activators (CRISPRa), repressors (CRISPRi), and epigenetic editors allow direct manipulation of local chromatin states to control gene expression (Chavez et al., 2015; Gilbert et al., 2013; Hilton et al., 2015; Liu et al., 2016). At the supranucleosomal scale, CRISPR-based imaging has enabled labeling and tracking of specific genomic loci in the 3-D nucleus of living cells through microscopy (Chen et al., 2013; Ma et al., 2016; Wang et al., 2019). Recently, several studies have developed hybrid approaches to combine CRISPR-based imaging with inducible dCas9-mediated direct manipulation of nuclear organization to investigate the consequences of these perturbations in real time. One study developed a system called CRISPR-Genome Organization (GO) that uses a small molecule inducible dCas9 to integrate targeted genomic loci into different nuclear compartments and measured the consequential changes to chromatin dynamics and transcription (Wang et al., 2018). The second study developed a system called CasDrop to optogenetically induce liquid-liquid phase separation at dCas9-bound telomeres to determine how chromatin responds to the formation of phase-separated droplets (Shin et al., 2018).

The combination of CRISPR-based imaging with targeted, inducible manipulation of chromatin composition likewise offers a promising approach to study the supranucleosomal organization and regulation of heterochromatin. In this study, we present CRISPR-Engineered Chromatin Organization (CRISPR-EChO), a platform for inducibly and reversibly tethering heterochromatin components across tens of kilobases of endogenous genomic regions. Using CRISPR-EChO, we investigated the supranucleosomal effects of tiling human HP1α across large genomic regions on chromatin state, gene expression, and dynamic chromatin behavior in live cells. We find that large-scale HP1α recruitment represses distal gene expression, promotes long-range interactions with other HP1α enriched regions, possesses structural stability, and compacts bound chromatin in a reversible manner. These findings substantiate the multi-faceted role of HP1α in shaping higher-order heterochromatin organization in eukaryotes.

## RESULTS

### Engineering and characterization of the CRISPR-EChO platform

To build the CRISPR-EChO system for synthetically recruiting heterochromatin components to targeted genomic regions in human cells, we adapted an abscisic acid (ABA)-inducible dCas9 protein design previously used for CRISPR-GO (Wang et al., 2018) and for dCas9-based transcriptional regulation (Gao et al., 2016) (**Fig. 1A**). We fused *S. pyogenes* dCas9 to the ABI domain of the ABA-induced ABI/PYL1 heterodimerizing system (Liang et al., 2011), along with a tagBFP marker for sorting and visualization. For the effector component, we fused the PYL1 domain of the ABA system to an sfGFP marker and the full length human HP1α protein. We also introduced an antibiotic-selectable single guide RNA (sgRNA) to target this protein module to a target site of interest. To generate large, highly concentrated HP1α domains visible by microscopy, we chose to target tandem repeat regions within the genome containing hundreds of copies of the binding site and spanning tens of kilobases of linear DNA (**Fig. 1B**).

**Figure 1.**
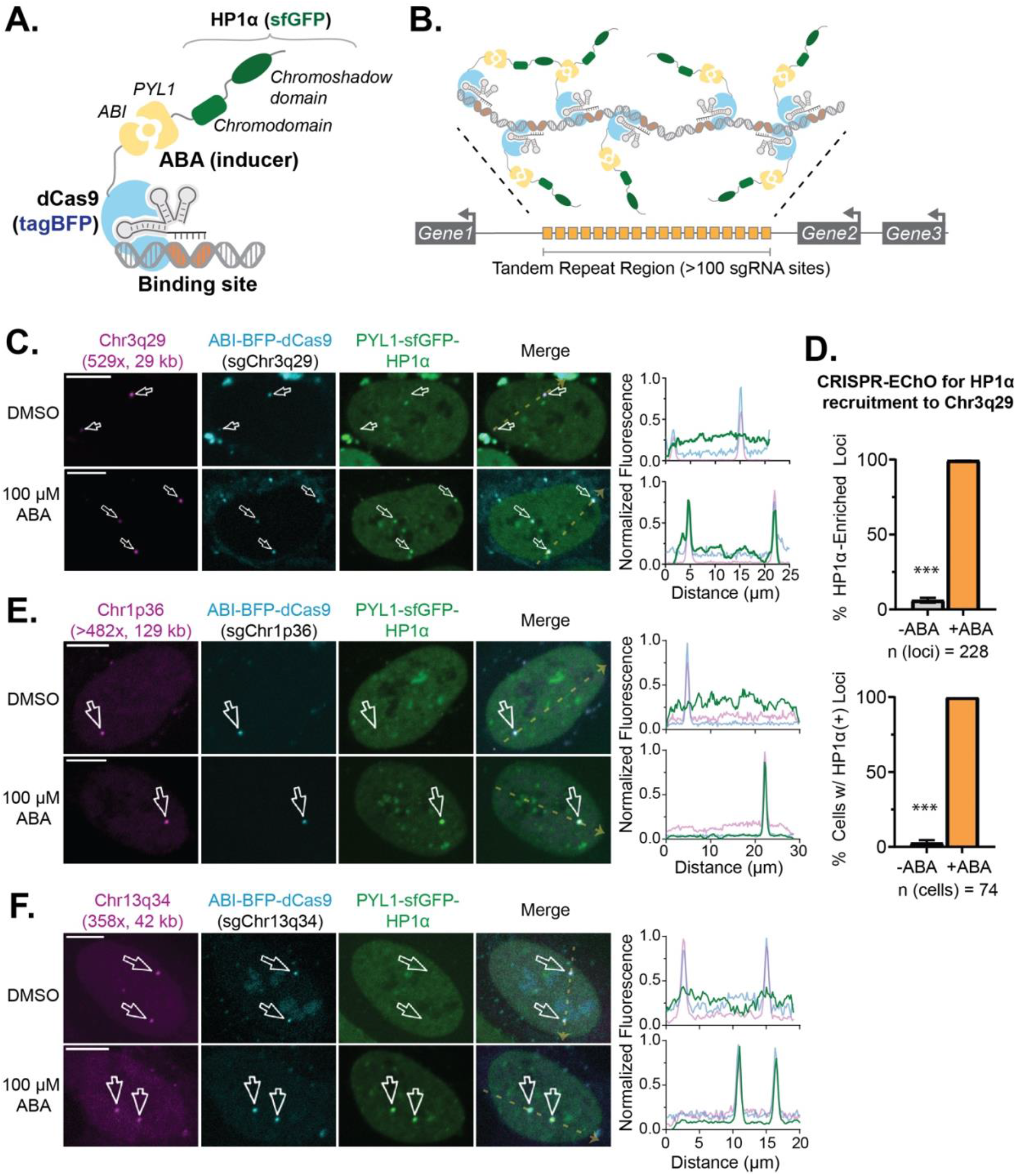
CRISPR-EChO Recruits HP1α En Masse at Endogenous Genomic Targets. **(A)** Schematic of the CRISPR-EChO protein design for recruiting full-length HP1α to target sites. The two structured domains of HP1α are labeled. Fused fluorescent protein markers are indicated but not depicted. **(B)** Schematic of the CRISPR-EChO strategy of tiling engineered proteins along large regions of genomic DNA. **(C)** Representative confocal microscopy images comparing localization of dCas9-HaloTag labeled Chr3q29, ABI-tagBFP-dCas9, and PYL1-sfGFP-HP1α in U2OS cells expressing the CRISPR-EChO system after 8 hours DMSO or 100 μM ABA treatment. **(D)** Quantification of PYL1-sfGFP-HP1α recruitment efficiency at labeled Chr3q29 loci before and after 1 hour 100 μM ABA addition. In the bottom panel, cells with >50% Chr3q29 loci with HP1α recruitment are considered HP1α(+). Grey bars indicate cells prior to ABA treatment, orange bars indicate cells after 1 hour ABA treatment. Error bars represent SEM calculated according to Bernoulli distributions. *** p < 0.001, two-sided Fisher’s exact test. **(E-F)** Representative confocal microscopy images showing recruitment of HP1α to alternative sgRNA-directed genomic tandem repeat sites on **(E)** Chr1p36 or **(F)** Chr13q34 after 8 hours DMSO or 100 μM ABA treatment. In **(C), (E)**, and **(F)**, magenta parenthetical text indicates total number of sgRNA binding sites and length of the repeat region in kilobases. White arrows indicate labeled loci, and right plots show the line scans along the yellow dotted line in each Merge image of fluorescence intensities for each color channel. Fluorescence intensities are normalized to the maximum (1) and minimum (0) intensities within the nucleus in the image. Scale bars represent 10 μm.

We stably expressed these components in human U2OS osteosarcoma cells through lentiviral transduction, followed by fluorescence activated cell sorting (FACS) and antibiotic selection. Visualization of the cell line via confocal microscopy showed that the ABI-tagBFP-dCas9 component in complex with an sgRNA targeting a 529-copy tandem repeat, 29 kilobase site on Chr3q29 successfully localized to the target site and formed discrete puncta (**Fig. 1C**). In the absence of the ABA inducer, PYL1-sfGFP-HP1α localized to natural heterochromatin within the nucleus (**Fig. 1C**), showing similar nuclear distribution as immunostained endogenous ortholog HP1β or lentivirally transduced ectopic mCherry-HP1α (**Fig. S1A-S1B**). Upon addition of 100 μM ABA, we observed recruitment of PYL1-sfGFP-HP1α to the dCas9-targeted site to form bright, rounded puncta (**Fig. 1C**). This recruitment was highly efficient, as one hour after inducer treatment the co-localization of PYL1-sfGFP-HP1α with Chr3q29 increased from 6.1% to 99.6% of all Chr3q29 loci, and the frequency of cells with >50% HP1α co-localized Chr3q29 loci increased from 2.7% to 100% (**Fig. 1D**). We also showed that sgRNAs targeted to other high copy tandem repeat sites across the genome enabled PYL1-sfGFP-HP1α recruitment to different chromosomal targets (**Fig. 1E**). These results demonstrate that CRISPR-EChO can inducibly and efficiently recruit a synthetic heterochromatin effector to targeted endogenous loci across different genomic contexts to form supranucleosomal scale structures.

### HP1α alone cannot fully convert a large targeted region to into heterochromatin

Natural HP1-associated heterochromatin is characterized by the enrichment of HP1 family proteins and repressive transcription factors such as KAP1 (TRIM28/TIFβ) and is epigenetically marked by H3K9me3 (Iyengar and Farnham, 2011; Maison and Almouzni, 2004). We next assessed to what extent synthetic HP1α tethering to target loci by the CRISPR-EChO system reproduces these hallmarks of natural heterochromatin. In U2OS cells expressing the CRISPR-EChO system, we transduced ectopic mCherry-HP1α as a proxy indicator for free-floating endogenous HP1α and immunostained for H3K9me3 and KAP1 to look for enrichment of these markers at the target site by confocal microscopy. First, in the absence of inducer, we found that the Chr3q29 site was not naturally heterochromatic, with low HP1α, H3K9me3, and KAP1 abundance relative to the signal across the nucleus (**Fig. S1C-S1E**). ABA addition to induce PYL1-sfGFP-HP1α localization to the Chr3q29 locus strongly increased mCherry-HP1α signal at the target site, suggesting that synthetic HP1α tethered through CRISPR-EChO can further recruit endogenous HP1 proteins (**Fig. 2A**). However, we did not observe notable enrichment of H3K9me3 or KAP1 at the target region (**Fig. 2A**). This finding was corroborated via ChIP-seq of H3K9me3 in cells with or without ABA addition. Addition of ABA to recruit HP1α did not alter H3K9me3 levels in regions surrounding the targeted tandem repeat or more broadly across Chr3q29 (**Fig. 2B, Fig. S2A**). These results raised the question of whether tethering HP1α alone is insufficient to produce these changes in a region of active euchromatin that may counteract heterochromatin formation.

**Figure 2.**
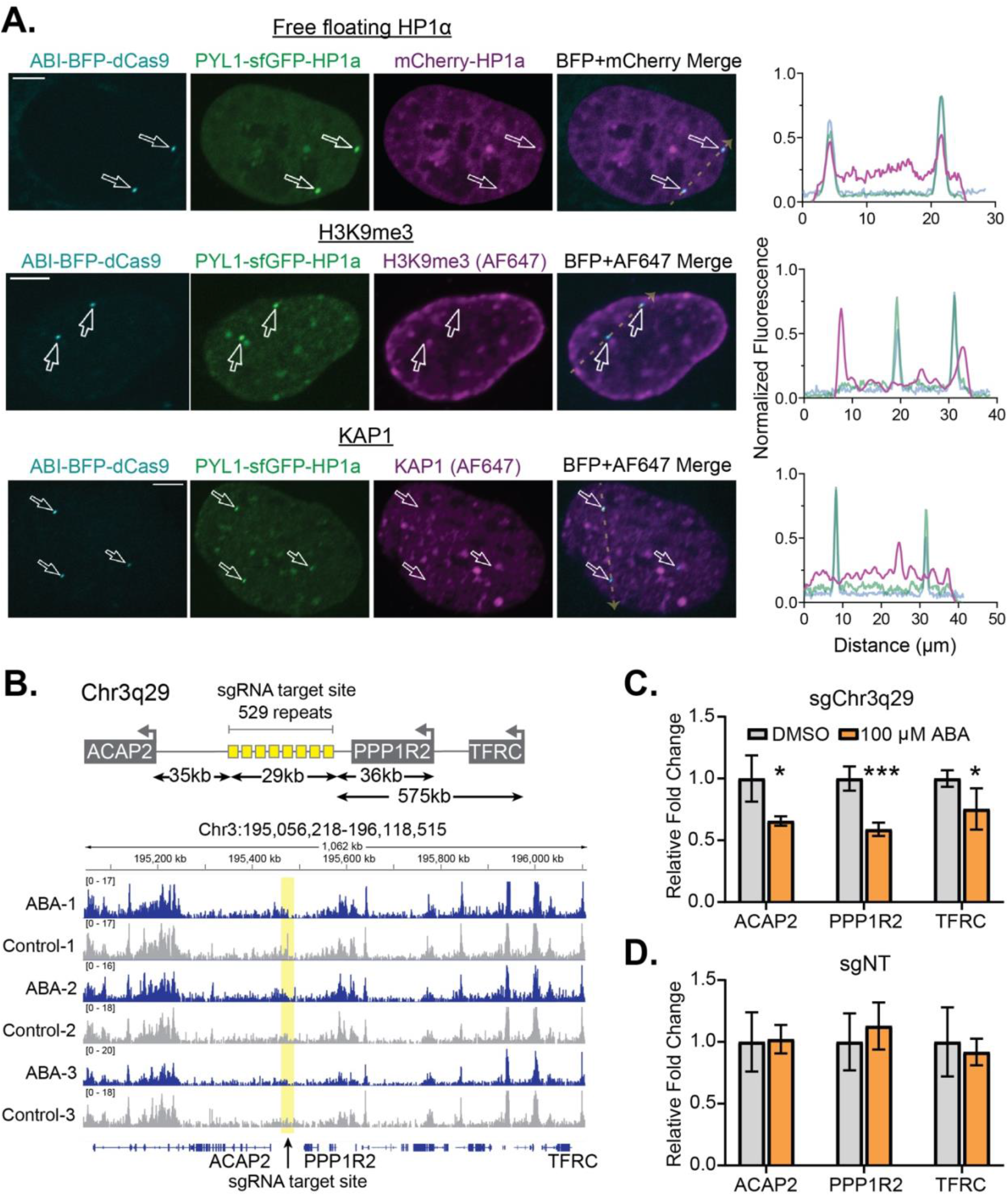
Heterochromatic Properties of Large-Scale Tethered HP1α at Chr3q29. **(A)** Representative confocal microscopy images of U2OS cells expressing the HP1α CRISPR-EChO system targeting Chr3q29 and treated with 100 μM ABA for two days. Top panel shows cells co-expressing mCherry-HP1α to represent the nuclear localization of free-floating (non-PYL1-fused) HP1α. Middle panel shows cells immunostained for H3K9me3 and visualized using an AlexaFluor 647-conjugated secondary. Bottom panel shows cells similarly immunostained for KAP1. Right plots show the line scans along the yellow dotted line in each Merge image of fluorescence intensities for each color channel. Fluorescence intensities are normalized to the maximum (1) and minimum (0) intensities within the nucleus in the image. Scale bars represent 10 μm. **(B)** H3K9me3 ChIP-seq IGV read coverage histogram tracks of a 1 Mb region of Chr3q29 surrounding the sgRNA target site for three replicate samples of CRISPR-EChO U2OS cells targeting Chr3q29 and treated with 100 μM ABA or DMSO control for two days. Track heights are normalized to total mapped reads for each sample. An illustration of the Chr3q29 locus and relative positions of the targeted tandem repeat site and genes *ACAP2*, *PPP1R2*, and *TFRC* is depicted on the left. Distances are measured from the edge of the tandem repeat site to the transcriptional start site of the respective gene. **(C-D)** RT-qPCR quantifying mRNA levels of the three indicated genes in U2OS cells expressing the HP1α CRISPR-EChO system and either **(C)** sgChr3q29 or **(D)** a non-targeting control sgRNA. Cells were treated with DMSO or 100 μM ABA for 5 days before RNA extraction. Data represents mean ± SD for four biological replicates from two independent experiments. * p < 0.05, *** p < 0.001, two-sided t-test.

### Large-scale HP1α recruitment represses distal gene expression

Heterochromatin is associated with gene repression, and numerous studies have shown that HP1α recruitment to gene promoters can repress expression of adjacent genes (Braun et al., 2017; Danzer and Wallrath, 2004; Hathaway et al., 2012; Li et al., 2003; Seum et al., 2001). We wanted to determine whether tethering of heterochromatin components by CRISPR-EChO functionally alters gene expression in large areas surrounding the targeted region. Using RT-qPCR, we quantified the expression of three highly expressed genes at Chr3q29 whose transcriptional start sites reside 35 kb (*ACAP2*), 36 kb (*PPP1R2*) and 575 kb (*TFRC*) away from the targeted tandem repeat site (**Fig. 2B**). Interestingly, we observed a moderate but significant repression of each of these three distal genes upon ABA-induced HP1α recruitment (**Fig. 2C**). When a non-targeting sgRNA control was used, ABA addition did not alter expression of any of the three genes (**Fig. 2D**). Looking by ChIP-seq, the repressive effect was not associated with corresponding changes in H3K9me3 at the respective gene promoter in ABA-treated cells (**Fig. S2B**). Introducing an I165E mutation to HP1α rendering it unable to dimerize (Norwood et al., 2006) abolished the repressive effect (**Fig. S2C**). Likewise, a truncated HP1α(Δ1-72) without the N-terminus and chromodomain required for oligomerization (Larson et al., 2017), but previously reported to remain competent for proximal gene repression (Braun et al., 2017; Hathaway et al., 2012), similarly abolished this distal repressive effect (**Fig. S2D**). These results suggest that the distal repressive capacity of HP1α requires a fully-intact protein that allows both dimerization and oligomerization and utilizes a repressive mechanism not reliant on H3K9me3, distinct from that previously reported at HP1α-tethered gene promoters.

### HP1α assembly can occur *de novo* or through fusion to existing heterochromatin

Given that CRISPR-EChO activity can be regulated by inducer addition to provide temporal control over HP1α recruitment, we sought to understand the dynamics of this process using time lapse microscopy. Since ABI-tagBFP-dCas9 was prone to photobleaching and phototoxicity upon repeated imaging, we modified our sgRNA system to include two MS2 hairpins (Konermann et al., 2014) and co-expressed MS2 coat protein (MCP)-mCherry to track the position of the target loci-bound dCas9 using mCherry instead of tagBFP (**Fig. 3A**). We observed that inducer addition caused synthetic HP1α puncta to rapidly localize to the target site and grow in a biphasic manner (**Fig. 3B-3C**). Puncta formed and grew quickly in the first 10 minutes after inducer addition, corresponding to recruitment of PYL1-sfGFP-HP1α through ABA-induced heterodimerization of ABI and PYL1. Subsequently, puncta entered a slow growth phase, which we speculate to correspond to additional recruitment of PYL1-sfGFP-HP1α through homodimerizing and oligomerizing interactions through HP1α (Larson and Narlikar, 2018). We also observed that there existed multiple modes of HP1α puncta nucleation at Chr3q29. While most puncta formed *de novo*, in the instances where Chr3q29 was positioned near or overlapping with pre-existing, chromocenter-like HP1α structures in the nucleus prior to inducer addition, synthetic HP1α recruited to Chr3q29 integrated the target locus into these pre-existing HP1α structures (**Fig. 3D, Fig. S3A-S3B**).

**Figure 3.**
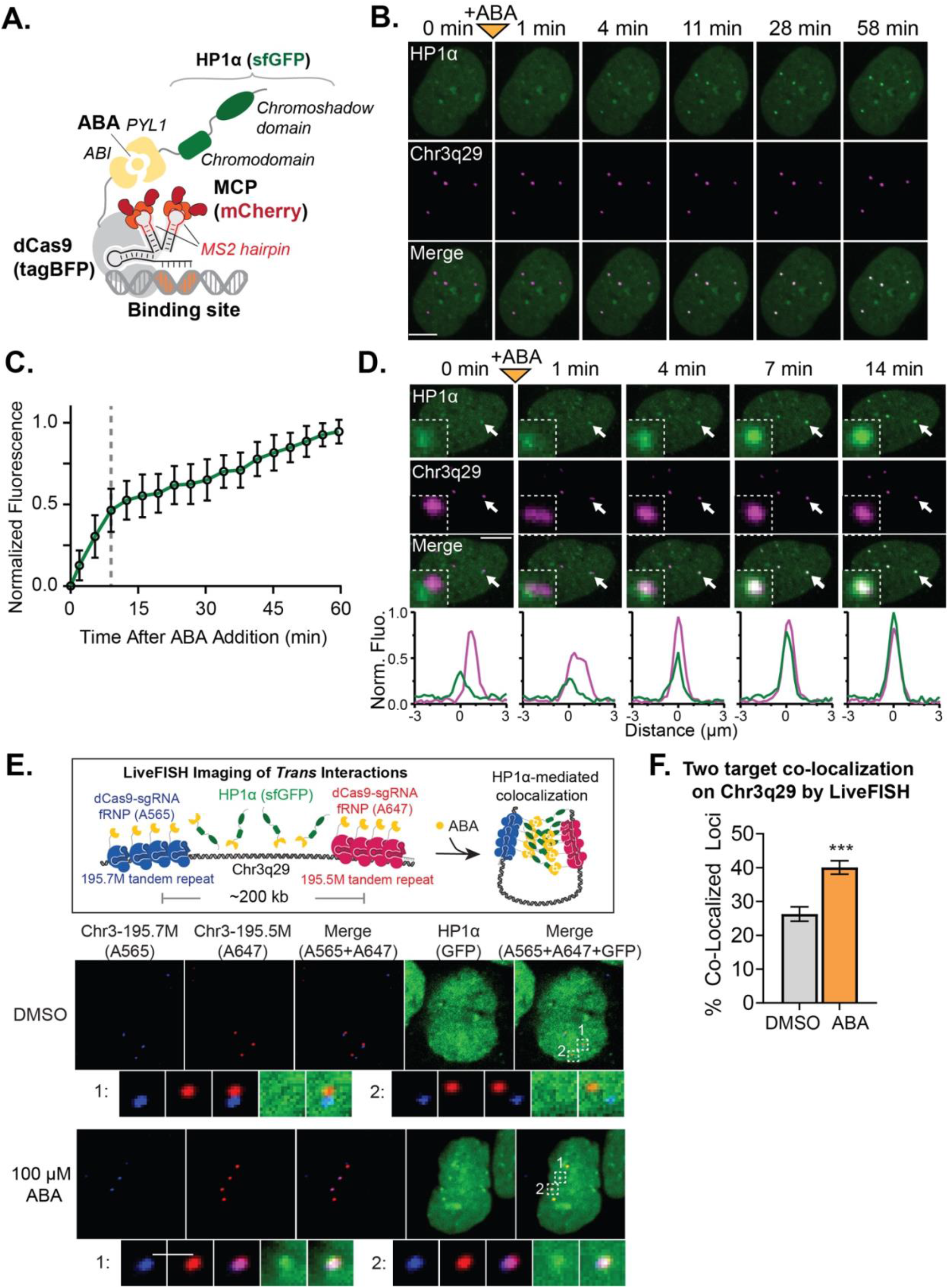
Dynamics of ABA-induced HP1α Assembly at Genomic Targets. **(A)** Illustration of the modified CRISPR-EChO system incorporating 2xMS2 hairpins and MCP-mCherry for improved target locus visualization. (**B**) Representative time-lapse confocal microscopy images of a U2OS cell expressing the HP1α CRISPR-EChO system showing *de novo* formation of HP1 heterochromatin at Chr3q29. Chr3q29-labeled images show location of MCP-mCherry bound to dCas9 at Chr3q29. Time 0 min represents image taken immediately before 100 μM ABA addition, and 1 min represents the first image taken of the same cell after addition. **(C)** Quantification of PYL1-sfGFP-HP1α signal intensity at Chr3q29 loci for time lapse microscopy. For each measured locus, the intensity before inducer addition is set to 0, and the maximum observed intensity is set at 1. The gray dotted line delineates the different phases of puncta growth. Data represent mean ± SD for 50 tracked loci from 12 cells. **(D)** Representative time-lapse confocal microscopy images showing incorporation of Chr3q29 into natural HP1α structures in the nucleus mediated by CRISPR-EChO. Time 0 min represents image taken immediately before 100 μM ABA addition, and 1 min represents the first image taken of the same cell after addition. Inset image represents 6x magnification of the region indicated by the white arrow. Bottom plots show line scans of fluorescence intensities for each color channel for region indicated by the white arrow. Fluorescence intensities are normalized to the maximum (1) and minimum (0) intensities observed across line scans for all displayed time points. Scale bars represent 10 μm. **(E)** Representative confocal microscopy images of LiveFISH experiment whereby two separate fluorescently-labeled tandem repeat regions on Chr3q29 (195.7M/Atto565 and 195.5M/Atto647) are targeted by CRISPR-EChO and treated with DMSO or 100 μM ABA for 6 hours. Bottom panels represent 4x magnification of the regions labeled 1 and 2 in the Merge image. Illustration of the LiveFISH experimental set-up. Atto565-labeled sgRNA targeting Chr3-195.7M and Atto647-labeled sgRNA targeting Chr3-195.5M tandem repeats on Chr3q29 are simultaneously introduced to U2OS cells expressing the CRISPR-EChO system are shown on the top. ABA addition leads to recruitment of HP1α to each tandem repeat and facilitates their interaction and co-localization with each other. **(F)** Quantification of the co-localization frequency of the Chr3q29-195.7M and Chr3q29-195.5M tandem repeat regions after 6 hours DMSO or 100 μM ABA treatment. Error bars represent SEM calculated according to Bernoulli distributions. *** p < 0.001, two-sided Fisher’s exact test. n=426 loci for the DMSO-treated group and 609 loci for ABA-treated group.

These observations suggested that HP1α can bridge heterochromatin to drive long-range chromatin interactions. To further explore this phenomenon, we used the LiveFISH method (Wang et al., 2019) to simultaneously introduce two fluorophore-conjugated sgRNAs targeting tandem repeat regions 200 kb apart on Chr3q29 into cells expressing the CRISPR-EChO system (**Fig. 3E-3F**). We observed that ABA-induced HP1α recruitment to these distal tandem repeat sites increased the frequency at which they co-localized from 26% to 40% (**Fig. 3F**). Altogether, these results indicate that HP1α can bring distal regions of chromatin together in 3-D space, likely due to its oligomeric properties. While tethered *Drosophila* HP1a has been reported to facilitate long-range loop formation in polytene chromosomes (Azzaz et al., 2014; Li et al., 2003; Seum et al., 2001), to our knowledge, similar behavior has not been previously observed in mammalian cells.

### HP1α puncta are resistant to disassembly

We performed similar time lapse microscopy to investigate the disassembly kinetics of PYL1-sfGFP-HP1α from the target site after inducer removal. Surprisingly, we observed that HP1α reduced the rate at which at which the sfGFP signal is lost at the target site. 60% of HP1α puncta at Chr3q29 remained at 1.5 hours after ABA washout, while 4.7% of puncta persisted for over 8 hours (**Fig. 4A-4B**). In contrast, for a control PYL1-sfGFP-NLS effector in which HP1α was removed and replaced by only an SV40 nuclear localization signal, only 2.0% of puncta remained after 1.5 hours, indicating that the ABI-PYL1 heterodimer normally dissociates rapidly when ABA is removed from the media (**Fig. 4B, Fig. S4A**). This HP1α conferred resistance to disassembly after ABA removal was also apparent at the Chr1p36 and Chr13q34 tandem repeat sites (**Fig. 4B, Fig. S4B**). Furthermore, we quantified the rate of loss for HP1α mutants with impaired dimerization/oligomerization at Chr3q29. The dimerization deficient I165E mutant showed a rate of loss nearly identical to that of the NLS control, while the oligomerization deficient truncated HP1α(Δ1-72) showed an intermediate rate of loss between that of HP1α and the NLS control effector (**Fig. 4B**). The difference in disassembly rates between HP1α, its mutants, and the NLS control effectors suggests that HP1α aggregates generated at target loci possess semi-stable structure that is resistant to disassembly, and that this stability is related to HP1α’s ability to multimerize.

**Figure 4.**
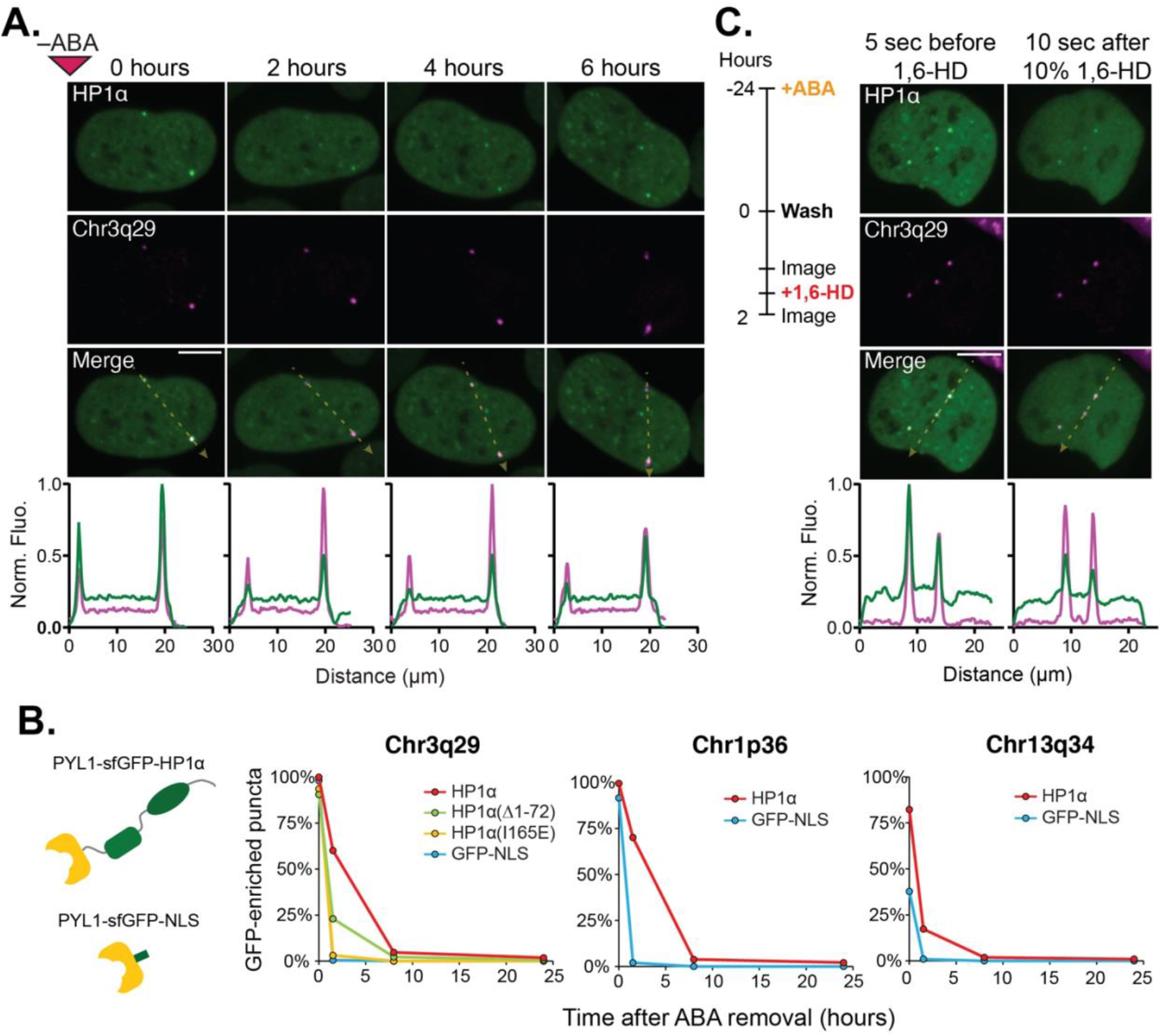
Assembled HP1α at Targeted Loci are Resistant to Disassembly. Representative time-lapse confocal microscopy images of a U2OS cell expressing the HP1α CRISPR-EChO system after ABA removal. Cells were treated for 24 hrs with 100 μM ABA prior to imaging. The 0 hour image represents the first image taken immediately after ABA removal. Bottom plots show line scans of fluorescence intensities for each color channel along the yellow dotted lines in each Merge image. Fluorescence intensities are normalized to the maximum (1) and minimum (0) intensities observed across line scans for all displayed time points. **(B)** Comparison of effector disassembly rates at Chr3q29, Chr1p36, and Chr13q34 between U2OS cells expressing CRISPR-EChO with PYL1-sfGFP-HP1α and a PYL1-sfGFP-NLS control, shown in schematic on left. For Chr3q29, disassembly rates of mutants PYL1-sfGFP-HP1α(Δ1-72) and PYL1-sfGFP-HP1α(I165E) were also analyzed. Graph displays the percentage of all indicated loci with GFP-enrichment at the indicated time points (n=128-234 loci). **(C)** Comparison of HP1α distribution at Chr3q29 loci and within the nucleus before and after 10% 1,6-hexanediol treatment. The experimental set-up is shown on the left, and representative confocal microscopy images are shown on the right. Bottom plots show fluorescence intensities of each color channel along the yellow dotted lines in each Merge image. Fluorescence intensities are normalized to the maximum and minimum (0) intensities observed across both line scans before and after treatment. Scale bars represent 10 μm.

We observed that a cell undergoing mitosis after ABA removal did not reform HP1α puncta at Chr3q29 in the daughter nuclei (**Fig. S5A**). We then asked whether disassembly of the HP1α puncta is dependent upon the active disruption of chromatin structures during DNA replication or mitosis. To test this hypothesis, we compared the disassembly rates between cells proliferating normally versus those arrested at the G1/S interface with hydroxyurea (**Fig. S5B-S5D**). G1/S-arrested cells did not exhibit changes in the disassembly rates of HP1α puncta compared to unarrested cells, indicating that the cell cycle is not a major determinant of the disassembly process.

Lastly, we tested whether disruption of multivalent HP1α interactions via 1,6-hexanediol (Strom et al., 2017) would disassemble semi-stable HP1α puncta at Chr3q29. Addition of 10% 1,6-hexanediol two hours after ABA removal globally disrupted HP1α nuclear structures (**Fig. 4C**). At Chr3q29, synthetic HP1α puncta showed strongly reduced signal intensity, though not complete HP1α loss (**Fig. 4C**). Since 1,6-hexanediol disrupts weak, multivalent HP1α interactions that allow polymerization, this result indicates that HP1α polymerization contributes to the disassembly resistance of HP1α structures after ABA removal. The remnant HP1α fraction after 1,6-hexanediol treatment may result from an inability to disrupt HP1α dimerization, incomplete disruption of multivalency, or a reduced rate of ABA loss from polymerized PYL1-sfGFP-HP1α.

### HP1α recruitment reversibly compacts local chromatin structure

*In vitro*, HP1α has been shown to compact nucleosomal arrays (Azzaz et al., 2014), and in CHO cells, HP1α-bound genomically amplified synthetic DNA arrays adopt a compact conformation (Verschure et al., 2005). We investigated whether HP1α recruitment to endogenous tandem repeat sites induced chromatin compaction. In the absence of inducer, 10.1% of MCP-mCherry labeled Chr3q29 loci exhibited an amorphous diffuse structure instead of a spherical compact structure, suggesting that some Chr3q29 loci are by default in an open chromatin state with a flexible conformation (**Fig. 5A**). When HP1α was recruited upon inducer addition, we observed the coalescence of the dynamic, diffuse Chr3q29 signal into a single strong, stable, and spherical signal (**Fig. 5B, Fig. S6A**). In some cases, the conformational change at Chr3q29 was concomitant with the initial appearance of the HP1α signal. However, in other instances, the coalescence event occurs minutes or hours after HP1α recruitment (**Movie 1**). The observed frequency of diffuse puncta decreased from 10.1% before ABA addition to 5.8% after one hour to 0.9% after 24 hours (**Fig. 5A**). In comparison, when the NLS control effector was used instead, no significant changes in diffuse puncta frequency are observed at Chr3q29 between the three time points (**Fig. 5A**). We observed similar compaction effects at tandem sites on Chr1p36 and Chr13q34, each of which showed reduced frequency of diffuse puncta after HP1α recruitment but not after NLS control effector recruitment (**Fig. 5A, Fig. S6B**). As with disassembly, we observe at Chr3q29 no compaction when a dimerization deficient HP1α(I165E) mutant is used and an intermediate effect when an oligomerization deficient HP1α(Δ1-72) mutant is used (**Fig. 5A, Fig. S6C-S6D**).

**Figure 5.**
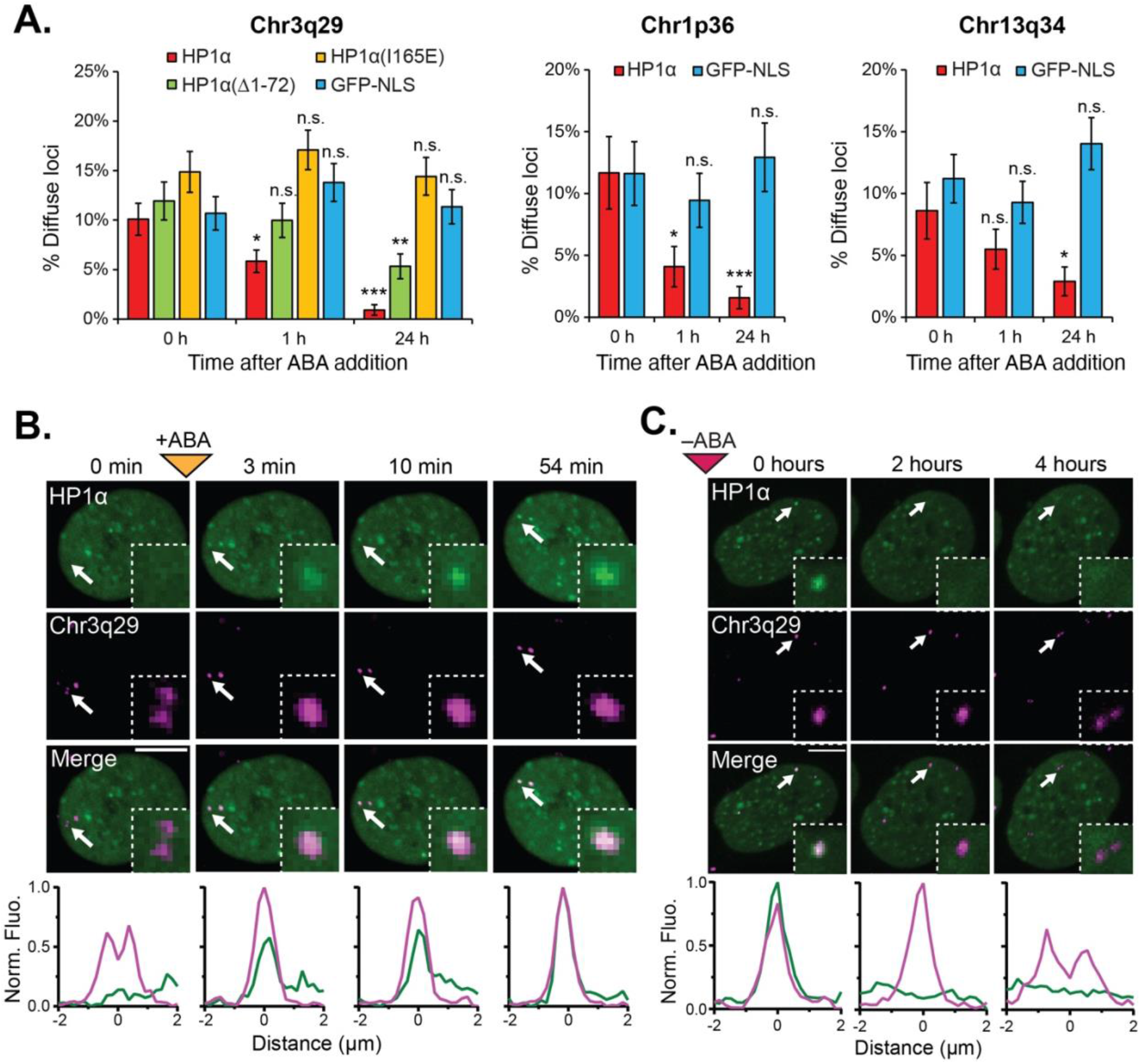
Inducible and Reversible Compaction of Endogenous Loci by CRISPR-EChO. **(A)** Quantification of percentage of Chr3q29, Chr1p36, or Chr13q34 loci displaying diffuse morphology in U2OS cells expressing CRISPR-EChO with the indicated effector domains before, after 1 hr, and after 24 hrs treatment with 100 μM ABA (n=285-428 loci). Error bars represent SEM calculated according to Bernoulli distributions. n.s. not significant, * p < 0.05, ** p < 0.01, *** p < 0.001, Fisher’s exact test between 0 h and indicated time points. **(B)** Representative time lapse confocal microscopy image showing compaction of a Chr3q29 locus upon 100 μM ABA addition, where compaction is concomitant with recruitment of the HP1α. Inset image represents 6x magnification of the region indicated by the white arrow. **(C)** Representative time lapse confocal microscopy image showing de-compaction of a Chr3q29 locus upon ABA removal. Decompaction occurs following loss of HP1α. Cells were treated with 100 μM for 24 hours prior to imaging. Inset image represents 4x magnification of the region indicated by the white arrow. In (**B-C**), bottom plots show fluorescence intensities of each color channel for region indicated by the white arrow. Fluorescence intensities are normalized to the maximum (1) and minimum (0) intensities observed across line scans for all displayed time points. Scale bars represent 10 μm.

Conversely, removal of ABA from the HP1α CRISPR-EChO cells enabled coalesced Chr3q29 loci to re-adopt a diffuse structure after HP1α was lost at the locus (**Fig. 5C**). These observations indicate that HP1α recruitment alone is sufficient to compact chromatin at a supranucleosomal scale and demonstrates that our synthetic biology approach can be used to dynamically manipulate chromatin conformation.

## DISCUSSION

### CRISPR-EChO: A synthetic biology platform for studying chromatin organization

In this work, we developed CRISPR-EChO, a CRISPR/dCas9-based approach to synthetically recruit and tile heterochromatin components across large regions of genomic DNA in a programmable and targeted manner. Combining CRISPR-based imaging with supranucleosomal scale perturbations to chromatin structure, CRISPR-EChO provides key advantages over previously developed methods for tethering heterochromatin components at single euchromatic sites. First, the inducibility and reversibility of the small molecule inducer system, combined with real-time visualization by microscopy, allows the study of temporal dynamics of alterations to chromatin state. Second, recruitment of heterochromatin components simultaneously to a region of the genome spanning tens of kilobases may produce emergent properties unobservable when targeting a single binding site such as a gene promoter. Lastly, a microscopy-driven approach offers inherent single cell readouts of chromatin organization. Such information can be combined with conventional epigenetic and transcriptional approaches to provide complementary information.

### HP1α recruitment creates novel chromosomal associations *in cis* and *in trans*

In our microscopy experiments, we observed two seemingly unrelated phenomena. First, HP1α recruitment to Chr3q29 can mediate long range chromatin associations in 3-D space, either by integrating a single target locus into pre-existing HP1α aggregates in the nucleus (**Fig. 3D, Fig. S3B**) or by integrating two target loci with each other (**Fig. 3E-3F**). Second, HP1α recruitment to Chr3q29 can cause an initially diffuse and unstructured genomic region to reversibly compact into a stable spherical conformation (**Fig. 5**). We believe that these two phenomena represent two consequences of the same property of HP1α: its ability to form dimeric, oligomeric, and polymeric assemblies. This property of HP1α has previously been demonstrated *in vitro* and its existence in cells has been hypothesized (Azzaz et al., 2014; Larson and Narlikar, 2018; Larson et al., 2017). In this context, the compaction and the long-range integration properties seem to demonstrate the ability of HP1α self-association to create *cis* and *trans* interactions, respectively. Compaction would result from HP1α bound to the far ends of the target site engaging in direct or indirect binding with one another, bringing the two ends into contact and maintaining a stable structure. This phenomenon has previously been reported in CHO cells, where EGFP-HP1α-lacR bound to a genomically-amplified synthetic LacO array exhibited a spherical structure while EGFP-lacR alone exhibited a fibrillar structure (Verschure et al., 2005). Here, we show that similar compaction can occur inducibly and reversibly at endogenous genomic sites upon HP1α recruitment.

The long range *trans* interactions of target-tethered HP1α with other HP1α-enriched regions, either natural or synthetic in origin, suggest that HP1α assemblies have a causal role in shaping of supranucleosomal-scale chromatin conformation. In support of this idea, in *Drosphila*, artificial HP1 binding sites on polytene chromosomes have been reported to generate novel chromosomal loops and contacts (Azzaz et al., 2014; Li et al., 2003; Seum et al., 2001). In the present study, we show, for the first time, the two processes of *cis* and *trans* associations occurring in real time at a human endogenous genomic locus upon HP1α recruitment. These supranucleosomal forms of HP1α self-association may be important factors in shaping the general spatial segregation of heterochromatin and euchromatin detected by Hi-C (Lieberman-Aiden et al., 2009; Rao et al., 2014) and observed by microscopy (Politz et al., 2013) to occur in the 3-D nucleus. Application of high-throughput conformation capture technologies, especially at the single-cell level, could reveal the extent to which HP1α associations alter the native chromatin conformation and could identify the locations of novel *trans* contacts.

### A potential passive mechanism for HP1α repression of distal gene transcription

In this study, HP1α was shown to be capable of repressing distal gene expression at Chr3q29 when recruited in large quantities (**Fig. 2C**). We posit that this repressive effect is the result of passive exclusion of RNA polymerase and transcriptional machinery due to steric effects of HP1α polymerization rather than the active epigenetic silencing of the assayed genes. The distinction between active and passive gene repression by HP1α has been proposed previously and may represent divergent consequences of HP1 binding to histone modification-sensitive regulatory elements in euchromatin versus polymerization-permissive, highly repetitive regions in constitutive heterochromatin (Eissenberg and Elgin, 2014; Hediger and Gasser, 2006). We present two lines of evidence in support for a passive mechanism of repression at Chr3q29.

First, despite impairing gene expression across Chr3q29, HP1α did not notably alter overall H3K9me3 levels across Chr3q29 as assayed by immunofluorescence (**Fig. 2A**) and ChIP-seq (**Fig. 2B**, **Fig. S2A**) or at the promoters of the three repressed genes (**Fig. S2B**). The lack of change in H3K9me3 suggests that the observed repression is not mediated by HP1α-induced epigenetic changes.

Second, the distal repression observed at Chr3q29 was also lost when full-length HP1α was replaced by a truncated Δ1-72 mutant lacking the N-terminus end and chromodomain (**Fig. S2D**). It has previously been shown that this chromoshadow mutant retains the histone modifying properties of HP1α and can silence gene expression when bound to promoters (Braun et al., 2017; Hathaway et al., 2012). However, the N-terminus and chromodomains of HP1α are essential for its ability to form oligomers (Canzio et al., 2013; Larson et al., 2017; Yamada et al., 1999). Therefore, the difference in distal repressive capacity at Chr3q29 between full-length HP1α and the mutant may be tied to these functions.

### HP1α polymerization confers resistance to dissociation from chromatin

We observed that HP1α forms semi-stable assemblies at the target locus, persisting long after ABA is depleted from the cell (**Fig. 4A-4B**). Treatment with 10% 1,6-hexanediol after ABA removal globally disrupted HP1α structures in the nucleus and significantly reduced HP1α occupancy of the Chr3q29 locus (**Fig. 4C**). The reduction in occupancy indicates that at least a fraction of bound HP1α at Chr3q29 is indeed present through weak, multivalent interactions consistent with HP1α polymerization. Furthermore, HP1α mutants in which dimerization or oligomerization function is eliminated showed either impaired or complete loss of this resistance to disassembly. These results suggest that HP1α polymers confer some degree of structural stability. The functional significance of HP1α polymer stability warrants further investigation. One intriguing possibility is that HP1α polymer stability may function as a buffer against stochastic nuclear perturbations to maintain constitutive heterochromatin in the compact state. Such a buffering effect would prevent catastrophic exposure of repetitive elements in the pericentromeres, which could lead to recombination and aberrant chromosomal segregation, or of telomeres, which could be recognized as double stranded breaks and lead to cell cycle arrest (Wallrath et al., 2014).

### A polymer model for HP1α regulation of chromatin organization

Collectively, these results support an HP1α polymer formation model to explain the various properties of CRISPR-EChO HP1α tethering to large genomic regions (**Fig. 6**). HP1α molecules tethered via ABA to dCas9 and tiled across a broad region of DNA engage in multivalent interactions with each other, with free-floating HP1α, and with HP1α bound to natural heterochromatin. These multivalent interactions between HP1α molecules create new contacts between different regions of chromatin and compacts chromatin structure in a manner that is reversible if HP1α is released from dCas9. The resulting HP1α polymer hinders access by transcriptional machinery to genes residing within the polymer and possesses a semi-stable structure resistance to disassembly. Finally, polymer formation may be dependent on the initial HP1α seeding concentration, such that it can only form when tiled across linear DNA in high-copy tandem binding sites in synthetic contexts or across large heterochromatin regions in natural contexts.

**Figure 6.**
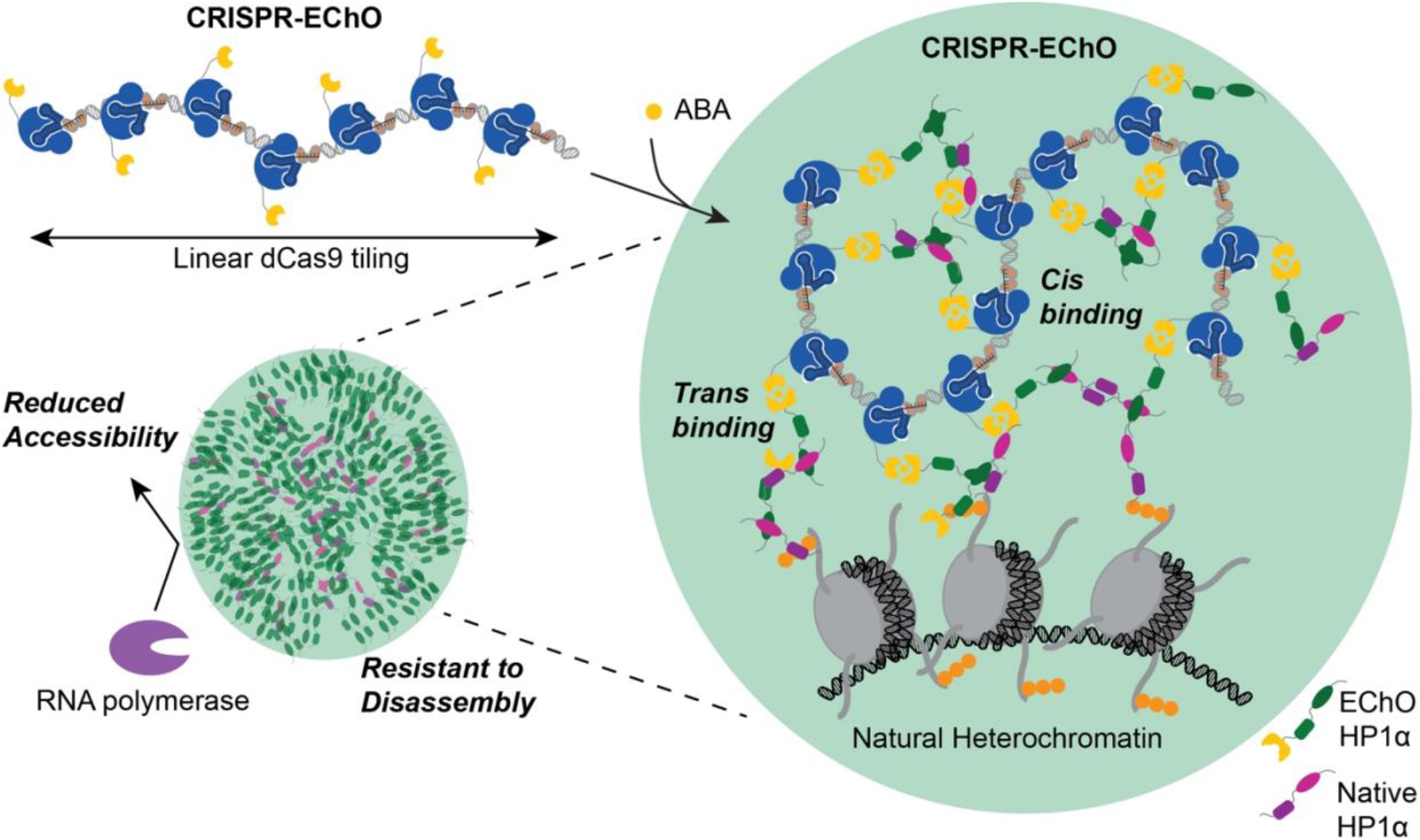
Polymer Model for HP1α Regulation of Chromatin Organization. In the absence of ABA inducer, the dCas9-bound tandem repeat region adopts a flexible and open conformation. Introduction of ABA induces dCas9 heterodimerization with engineered HP1α (green). Additional engineered HP1α or endogenous HP1α (magenta) are recruited to the site via HP1α homodimerization and oligomerization. HP1α generates *cis* interactions with other sites along the tandem repeat region and *trans* interactions with HP1α at other loci, either within natural heterochromatin or at additional dCas9-bound distal sites. These interactions lead to compaction of the target locus, co-localization of multiple targeted loci, and integration with natural heterochromatin. Collectively, the recruited HP1α forms a polymer that reduces accessibility of the encapsulated loci by transcriptional machinery and confers a semi-stable structure with delayed disassembly kinetics upon ABA removal.

### Phase separation of chromatin-bound synthetic HP1α assemblies

A remaining question of interest is whether the synthetic HP1α assemblies formed in a targeted manner undergo liquid-liquid phase separation. There is increasing evidence that liquid-liquid phase separation is a major driver of nuclear compartmentalization and that chromatin is subject to various forms of liquid-liquid phase separation (Strom and Brangwynne, 2019). Some HP1 orthologs, including human HP1α, have likewise been shown to undergo liquid-liquid phase separation *in vitro* and in cells due to the multivalency of its intrinsically disordered N-terminus and hinge regions (Larson et al., 2017; Sanulli et al., 2019; Strom et al., 2017). Several observations of HP1α made in this study are consistent with liquid-liquid phase separated behavior. First, HP1α recruited to target regions form spherical structures and can undergo fusion with other natural HP1α assemblies. Second, the aliphatic alcohol 1,6-hexanediol can partially disrupt these structures. Third, HP1α-bound chromatin appears to be less accessible by proteins like RNA polymerases. However, these behaviors could also be possible through interactions of non-phase separated HP1 multimeric assemblies. Indeed, a recent study found that mouse HP1α polymers at heterochromatin foci in living cells share the same phase as surrounding euchromatin despite phase separating *in vitro* at high concentrations (Erdel et al., 2020). Experimental determination of liquid-liquid phase separation remains challenging in cells (Alberti et al., 2019), and further exploration of this phenomenon for synthetic, chromatin-bound HP1α assemblies is warranted.

## Conclusions

Over the past 8 years, CRISPR/Cas9-based genome editing and transcriptional regulation have matured into established techniques to fundamentally change how biological research is conducted. More recently, CRISPR/Cas9-based epigenetic editing is poised to do the same. The development of CRISPR/Cas9 technologies for studying higher-order nuclear organization remains in its infancy. In the past two years, we have seen the first proof-of-concept studies demonstrating its potential in cells to dissect the complexity of the nucleus, including those that induce chromatin-bound liquid-liquid phase separation (Shin et al., 2018), tether loci to specific nuclear compartments (Wang et al., 2018), and, in the current study, tile heterochromatin components across large genomic regions. We envision a new era of dCas9-based tools to perturb and dissect the dynamic large-scale 3-D organization of chromatin in living cells in the future.

## ACKNOWLEDGEMENTS

The authors thank the members of the Qi lab for discussions and feedback and Aaron Gilter for helpful comments on the manuscript. We thank the Cell Sciences Imaging Facility at Stanford University for instrumentation used in this study. We acknowledge Luke Lavis at the Janelia Research Campus for providing the JF549-HaloTag ligand. L.S.Q. acknowledges support from the NIH Office of the Director (OD), the 4D Nucleome project, the NSF CAREER Award, and the Li Ka Shing Foundation. Y.G. acknowledges support from the Stanford Cancer Biology Graduate Program and the NSF GRFP fellowship. This work was supported by the NIH 4DN project under Grant No. 1U01DK127405, the National Science Foundation CAREER Award under Grant No. 2046650, and the Li Ka Shing Foundation.

## AUTHOR CONTRIBUTIONS

Y.G. and L.S.Q. conceived the research and designed the study. Y.G., M.H., and S.S. performed the experiments with assistance from H.W. Y.G., M.H., S.S., H.W. and L.S.Q analyzed the data. Y.G. wrote the manuscript and L.S.Q revised the manuscript with input from all authors.

## METHODS

### Cell Lines

U2OS (human bone osteosarcoma epithelial, female, ATCC HTB-96) and LentiX 293T (human embroyonic kidney epithelial, female, Takara 632180) cells were cultured in Dulbecco’s modified Eagle medium (DMEM) with GlutaMAX, 25 mM D-Glucose, and 1.0 mM sodium pyruvate (Life Technologies 10569044) supplemented with 10% FBS (Gibco 26140079) in incubators at 37°C and 5% CO_2_. Cell lines were not authenticated after purchase prior to use.

### Plasmid construction

Individual constructs for dCas9s, effectors and sgRNAs used in this study are described in **Table S1**. Nucleotide sequences for sgRNAs are provided in **Table S2**. New constructs from this study will be available on Addgene.

#### dCas9 fusions

Human codon-optimized optimized *Streptococcus pyogenes* dCas9 with two C-terminal SV40 NLSs was fused at the N-terminus to the ABI domain (gift from Jerry Crabtree, Addgene plasmid #38247) and tagBFP. Expression of the dCas9 was driven by a PGK promoter on a pHR backbone.

#### Chromatin effector fusions

Individual effectors were fused at the N-terminus to the PYL1 domain (gift from Jerry Crabtree, Addgene plasmid #38247) and sfGFP. Expression of the effector was driven by a PGK promoter on a pHR backbone.

HP1α, HP1α(I165E), and HP1α(Δ1-72) were cloned from GFP-HP1α (gift from Tom Misteli, Addgene plasmid #17652

#### sgRNAs

sgRNA expression was driven by a murine U6 promoter on a pHR backbone containing a CMV promoter-driven puromycin resistance gene. Modified sgRNA(2xMS2) expression was driven by a murine U6 promoter on a pHR backbone containing a UbC promoter-driven MCP-mCherry-p2A-puromycin resistance gene. MCP was cloned from pJZC116 (Addgene plasmid #62344).

#### Fluorescent HP1α reporter

The full-length human HP1α protein was fused at the N-terminus to mCherry. Expression was driven by a PGK promoter on a pHR backbone

### Lentivirus production

Lentivirus was generated by transiently transfecting LentiX 293T cells in 10 cm plates with the pHR construct of interest, pCMV-dR8.91, and pMD2.G at a ratio of 9:8:1 by mass using TransIT-LT1 transfection reagent (Mirus MIR 2306) at a ratio of 3 μL transfection reagent per 1 μg plasmid. Cell media was changed 24 h after transfection. Viral supernatant was collected 48 h post-transfection, passed through a 0.45 μm PVDF filter, and concentrated 10x using the Lentivirus Precipitation Solution (Alstem VC100) by centrifugation at 4°C for 30 m at 1,500x g following overnight incubation at 4°C.

### Cell line generation

To generate cell lines expressing the ABI-tagBFP-dCas9 construct, U2OS cells in 6-well plates were transduced at 50% confluence with 200 μL of 10x concentrated lentivirus and 5 μg/mL polybrene (EMD Millipore TR-1003-G). Approximately 4 days after transduction, cells were clonally sorted by FACS using a SONY SH800S sorter for tagBFP expression. Individual clones were expanded, and a low tagBFP expressing clone was selected as a base cell line.

PYL1-sfGFP-Effector constructs were similarly transduced into the base cell line and bulk sorted by FACS for sfGFP expression. PYL1-mCherry-Effector constructs were similarly transduced into the base cell line and bulk sorted by FACS for mCherry expression. Combinatorial effector cell lines were made by transducing PYL1-sfGFP-Effector constructs into ABI-tagBFP-dCas9 PYL1-mCherry-Effector cell lines.

For chromosomal locus imaging in **Figure 1**, a dCas9-HaloTag construct was similarly transduced into cells stably expressing dCas9 and effectors without sorting or selection.

For mCherry-HP1α imaging, an mCherry-HP1α construct was similarly transduced into cells stably expressing dCas9 and effectors without sorting or selection.

To introduce sgRNA to the above cell lines, cells in a 12-well plate were transduced at 50% confluence with 100 μL of 10x concentrated lentivirus and 5 μg/mL polybrene. 2 days after transduction, cells were selected for 3 days with 2 μg/mL puromycin.

### Imaging for chromosomal labeling/mCherry-HP1α

For chromosomal locus imaging in **Figure 1**, CRISPR imaging was used to visualize the location of the Chr3q29, Chr1p36, and Chr13q34 loci. Stable cell lines expressing CRISPR-EChO, dCas9-HaloTag, and sgRNA were seeded in 24-well μ-plate (Ibidi 82406) and treated with 100 μM abscisic acid (Sigma A1049) or DMSO (Sigma D2650) vehicle for 8 to 24 h. Cells were stained with JF549-HaloTag ligand (gift from Luke Lavis in Janelia Research Campus(Grimm et al., 2015)) at 10 nM for 15 m at 37°C in culture media. Cells were then washed twice with culture media, incubating for 30 m at 37°C during the second wash, then replaced with culture media containing fresh inducer before imaging.

For mCherry-HP1α imaging, U2OS cells stably expressing CRISPR-EChO and sgRNA were seeded at 10% confluence in 24-well μ-plate (Ibidi 82406) and treated with 100 μM abscisic acid or DMSO vehicle for 2 days before imaging.

### Immunostaining of heterochromatin markers

U2OS cells stably expressing CRISPR-EChO and sgRNA were seeded at 10% confluence in 24-well μ-plate (Ibidi 82406) and treated with 100 μM abscisic acid (Sigma A1049) or DMSO (Sigma D2650) vehicle for 2 days. Cells were fixed in 4% paraformaldehyde (ThermoFisher 28906) for 15 m, then rinsed 3 times with DPBS (Life Technologies 14190-250) for 5 m each. Fixed cells were blocked in 5% normal donkey serum (Jackson ImmunoResearch Laboratories 017-000-121) and 0.3% Triton X-100 (ThermoFisher 85111) in DPBS for 1 h at room temperature, then incubated overnight in at 4°C with primary antibodies. Samples were rinsed 3 times with DPBS for 5 m each, then incubated with secondary antibody 2 h at room temperature in the dark with agitation. Samples were rinsed 3 times with DPBS for 5 m each, then stored in DPBS for imaging.

Primary antibodies used in this study are Anti-Histone H3 (tri methyl K9) antibody - ChIP Grade (Abcam ab8898) used at 1:500 dilution, Anti-CBX1 / HP1 beta antibody (Abcam ab10478) used at 1:200 dilution, and Anti-KAP1 antibody - ChIP Grade (Abcam ab10483) used at 1:500 dilution. The secondary antibody used in this study is Alexa Fluor® 647 Donkey Anti-Rabbit IgG (H+L) Antibody (Life Technologies A31573) used at 1:500 dilution.

### ChIP-seq

Three clonal populations of U2OS cells expressing ABI-tagBFP-dCas9 and PYL1-sfGFP-HP1α with sgChr3 were grown to 80% confluency in 2×15-cm plates. Roughly 4 × 10^7^ cells per sample were washed with PBS before addition of 1% paraformaldehyde in fresh media for crosslinking. After 15 minutes, the samples were quenched with glycine to a final concentration of 0.125 mol/L glycine. Cells were then lifted with a cell scraper, spun down, and resuspended in SDS lysis buffer (1% SDS, 10 mmol/L EDTA, 50 mmol/L Tris-HCl, pH 8.1) at a ratio of 1 mL per 2 × 10^7^ cells and frozen at −80 before lysis at room temperature for 1h immediately before sonication. Pulse sonications were performed for 7 minutes at 40% amplitude with 1s on/1 s off on a Qsonica Q500 in 1mL aliquots in 1.5mL Eppendorf tubes with a total of 2 mL with maximum 50% SDS Lysis Buffer solution diluted with ChIP Dilution Buffer (0.01% SDS, 1.1% Triton X-100, 1.2 mmol/L EDTA, 16.7 mmol/L Tris-HCl, pH 8.1, 167 mmol/L NaCl). ChIPs were performed with antibodies for H3K9me3 (Abcam ab8898). Pulldowns were performed with Dynabeads Protein A (10001D, Lot 00531421). DNA quantities were measured by Qubit 2.0 (Invitrogen Qubit dsDNA HS Assay Kit, Q32854) and Bioanalyzer before sequencing with Novogene targeting 20M reads / sample.

Paired-end reads were acquired from Novaseq. Reads were aligned using bowtie2 (Langmead and Salzberg, 2012) and duplicates removed by samtools. Bigwig files were created by bamCoverage and imported into IGV for viewing. Genomic coverage by bedtools multicov was performed over 5kb bins created by bedtools makewindows.

### RT-qPCR

On day 0, U2OS cells stably expressing CRISPR-EChO and sgRNA were seeded at 12.5% confluence in 12-well plates and treated with 100 μM abscisic acid or DMSO vehicle. On day 3, cells were passaged at a 1:2 ratio and treated with fresh 100 μM abscisic acid or DMSO vehicle. On day 5, total RNA was extracted using the RNeasy Plus Mini Kit (Qiagen 74134), followed by cDNA synthesis using the iScript cDNA Synthesis Kit (BioRad 1708890). Quantitative PCR was performed using the SYBR Green Max Mix (BioRad 172-5125) with PrimeTime qPCR primers (IDT). Samples were run on a BioRad CFX384 Touch Real-Time PCR Detection System in technical duplicates for each experimental replicate. Gene expression was quantified using the 2^−ΔΔCT^ method (Livak and Schmittgen, 2001) using GAPDH as an internal control.

Primer sequences for qPCR and catalog numbers are provided in **Table S4**.

### ON and OFF time course experiments

For ON time course, on day 0, U2OS cells stably expressing CRISPR-EChO and sgRNA dissociated in TrypLE without phenol red (Life Technologies 12563-011) were seeded at 35% confluence in 96-well μ-plate (Ibidi 89626) in FluoroBrite™ DMEM (ThermoFisher A1896702) supplemented with 10% FBS and GlutaMAX (ThermoFisher 35050-061). On day 1, two hours before imaging, cells were treated with ProLong™ Live Antifade Reagent (ThermoFisher P36975). 100 μM abscisic acid was added to cells immediately after the Time 0 image is taken by adding a 3x concentrated stock to the imaging well.

For OFF time course, the same protocol was maintained with the following changes. 100 μM abscisic acid was added on Day 0 at time of seeding. On day 1, immediately before imaging, cells were washed 3 times with DPBS (Life Technologies 14190-250) then cultured in complete FluoroBrite DMEM media.

### Multi-locus labeling with LiveFISH

The unlabeled Cas9 tracrRNAs with an A-U flip in the original Cas9 cr:tracrRNA backbone sequence (Chen et al., 2013; Wang et al., 2019), Cas9 crRNAs targeting Chr3-195.7M labeled with Atto565 at the 5’ end, and Cas9 crRNAs targeting Chr3-195.5M labeled with Atto647 at the 5’ end were all synthesized by IDT. Nucleotide sequencers for spacers used for crRNAs are provided in **Table S3**. To prepare Cas9 gRNAs for LiveFISH, tracrRNAs and fluorescently labeled crRNAs were mixed at equal molar ratio, annealed in folding buffer (20 mM HEPES, pH7.5, 150 mM KCl), incubated at 95°C for 5 min, 70°C for 5 min, gradually cooled down to room temperature and supplemented with 1 mM MgCl_2_, followed by incubation at 40°C for 5 min and gradually cooled down. Then the fluorescently labeled gRNA_Chr3-195.7M and gRNA_Chr3-195.5M were mixed at equal molar ratio. 10 pmol of the mixed gRNAs together with 0.25 μL electroporation enhancer (IDT, 100uM, Cat#1075915) were transfected into 4×10^5^ suspended U2OS pSLQ4356 L-A5 pSLQ6901 cells using the standard protocol for the Neon Transfection System 10 μL kit (Thermo Fisher, MPK1025). The transfected cells were plated in 96-well ibidi μ-plates and treated with 100 μM ABA or DMSO while seeding for 6 h. Then cells were fixed with 4% PFA and imaged on a Nikon TiE inverted spinning disk confocal microscope. The imaging conditions are 30% power and 1 s exposure for 561 nm channel, 30% power and 1 s exposure for 640 nm channel, 50% power and 500 ms exposure for 480 nm channel, and 11 Z stacks at 0.5 μm step size.

### Cell cycle arrest

On day 0, U2OS cells stably expressing CRISPR-EChO and sgRNA were seeded at 7.5% confluence in 24-well μ-plate (Ibidi 82406) in FluoroBrite™ DMEM (ThermoFisher A1896702) supplemented with 10% FBS and GlutaMAX (ThermoFisher 35050-061). On day 1, cells were serum starved by in FluoroBrite DMEM with 0.1% FBS and GlutaMAX. On day 2, cells were switched back to FluoroBrite DMEM with 10% FBS and GlutaMAX and treated with 4 mM freshly prepared hydroxyurea (Sigma H8627) and 100 μM abscisic acid (Sigma A1049). On day 3, cells were washed 2 times with DPBS to remove ABA, then cultured in FluoroBrite DMEM with 10% FBS, GlutaMAX, and 4 mM freshly prepared hydroxyurea immediately before imaging. Control cells were similarly treated but without hydroxyurea on days 2 and 3.

### Flow cytometry

To measure the efficiency of cell cycle arrest by hydroxyurea, U2OS cells using for imaging were stained with Hoechst 33342 (ThermoFisher H3570) at 10 μg/mL for 15 m at 37 °C, washed once with DPBS, then dissociated with 0.25% Trypsin-EDTA (Life Technologies 25200056) and neutralized with standard DMEM with GlutaMAX and 10% FBS. Cells were passed through a 45 μm strainer then immediately analyzed on a Beckman Coulter CytoFlex S using a 405 nm laser. Cell cycle analysis was performed using FlowJo (FlowJo, LLC).

### Multivalency disruption

U2OS cells stably expressing CRISPR-EChO and sgRNA were treated according to the OFF time course protocol above. Two hours after ABA removal, cells were treated with 10% w/v 1,6-hexanediol (Sigma 240117) by adding a 3× concentrated stock to the imaging well. Cells were imaged immediately before and immediately after hexanediol addition.

### Microscopy

Microscopy was performed on a Nikon TiE inverted spinning disk confocal microscope equipped with a Photometrics sCMOS Prime95B camera and 405 nm, 488 nm, 561 nm, and 642 nm lasers using a 60x PLAN APO oil objective (NA=1.40). Images were taken using NIS Elements version 4.60 software by time-lapse microscopy with Z stacks at 0.5 μm steps. Live cells were kept at 37°C and 5% CO_2_ during imaging.

### Image processing

Image processing was performed in FIJI (Schindelin et al., 2012). For immunofluorescence and mCherry/mIFP-HP1α enrichment analysis, a single Z plane showing maximum fluorescence of the labeled loci, or the maximum Z projection of two to three consecutive Z planes capturing maximum loci fluorescence are shown in figures. For time-lapse experiments, the maximum Z projection of 9 to 11 Z stacks at 0.5 μm are shown in figures and used for fluorescence intensity quantification. Registration by the Correct 3D Drift (Parslow et al., 2014) plugin in FIJI was used to re-align cells across frames of time.

Line scans were performed using the “Analyze/Plot Profile” function in FIJI on a line drawn to intersect two labeled genomic loci and extending outside the nuclear boundary in both directions. Fluorescence intensity for each channel was calculated at each point alone the line and normalized to the maximum (=1) and minimum (=0) fluorescence intensity values observed in the image within the cell nucleus. For time course experiments, fluorescence values were normalized to the maximum or minimum fluorescence intensities observed across all line scans across all displayed time points.

For the OFF time-lapse experiments, variations in overall GFP image intensities at different time points due to instrumentation were corrected using the Bleach Correction macro (J. Rietdorf, EMBL Heidelberg) with an ROI drawn around regions of the nucleus excluding synthetic puncta at target loci prior to analysis.

### Quantification of effector enrichment

For quantification of effector recruitment efficiency, target loci were labeled with MCP-mCherry, which binds to the sgRNA(2xMS2). Enrichment of GFP at each labeled locus was determined on an individual Z plane basis for a Z stack image without double counting. Loci were binned into two categories: loci that show GFP enrichment and loci that do not show GFP enrichment. Enrichment was determined by performing two perpendicular line scans through each MCP-mCherry-labeled locus, where enriched loci show a quantitative increase in GFP signal intensity coincident with the mCherry peak compared to the GFP signal ∼2 μm (approximate mCherry peak diameter) in each direction for both lines. The number of loci for each category was recorded for individual cell. Cells were then binned into cells showing >50% loci with GFP enrichment (positive) and cells showing ≤50% loci with GFP enrichment (negative).

For quantification of GFP fluorescence intensity over time shown in **Figure 3**, the TrackMate (Tinevez et al., 2017) plugin in FIJI was used to automatically identify and track particles. Individual fluorescence measurements were calculated as mean fluorescence intensity within particles of 1 μm diameter using TrackMate. For early time points in which no particles have formed at target loci, the mean fluorescence intensity was manually determined using the “Analyze/Measure” function in FIJI based on the location of the labeled locus in the mCherry channel. For normalization, fluorescence intensity at time 0 was set to 0, and the maximum fluorescence intensity observed during the time course was set to 1.

### Determination of locus conformation

For binary determination of whether a locus was in a diffuse state or compact state, a line scan was performed through its major (longest) axis within a single Z-plane. A locus was defined as compact if the mCherry signal intensity along the axis showed a single sharp, distinct peak or diffuse if the mCherry signal intensity showed multiple peaks or no distinct peaks.

### Statistical Analysis

For quantification of effector enrichment, *p* values were calculated using two-sided Fisher’s exact test in Prism 8 (GraphPad), and error bars show the standard error of the mean (SEM) calculated according to Bernoulli distributions. The number of counted loci and cells are listed for each figure in the legends.

For gene expression measurements, *p* values were calculated using two-sided t-test without assuming equal variance in Prism 8 (GraphPad), and error bars show the standard deviation. The number of experimental replicates is listed for each figure in the legends.

---

**Figure S1.**
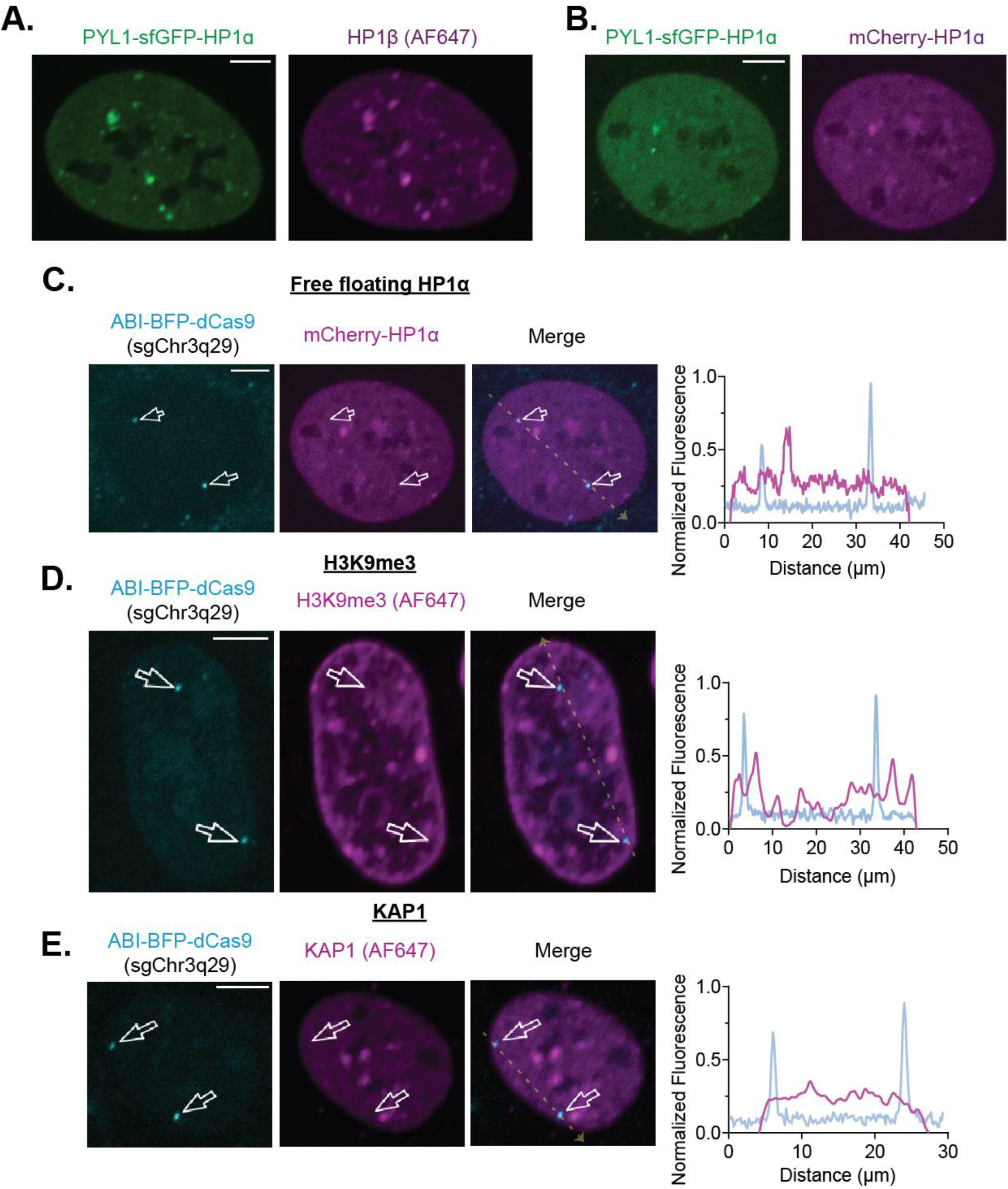
Heterochromatic Characterization of CRISPR-EChO Control U2OS Cells. **(A)** Representative confocal microscopy image showing PYL1-sfGFP-HP1α with similar nuclear localization as endogenous HP1β. U2OS cells expressing PYL1-sfGFP-HP1α are immunostained for HP1β and visualized using an AlexaFluor 647-conjugated secondary. **(B)** Representative confocal microscopy image showing PYL1-sfGFP-HP1α with similar nuclear localization as ectopic mCherry-labeled HP1α. **(C)** Representative confocal microscopy image showing that ABI-tagBFP-dCas9 bound Chr3q29 loci do not exhibit high mCherry-HP1α signal. The same cell is visualized in **(B)** and **(C)** for different purposes. **(D)** Representative confocal microscopy image showing that ABI-tagBFP-dCas9 bound Chr3q29 loci do not exhibit high H3K9me3 signal. Cells were immunostained for H3K9me3 and visualized with an AlexaFluor 647-conjugated secondary. **(E)** Representative confocal microscopy image showing that ABI-tagBFP-dCas9 bound Chr3q29 loci do not exhibit high KAP1 signal. Cells were immunostained for KAP1 and visualized with an AlexFluor 647-conjugated secondary. In **(C-E)**, white arrows indicate nuclear positions of Chr3q29. Right plots show line scans of fluorescence intensities for each color channel along the dotted yellow line in each left image. Fluorescence intensities are normalized to the maximum (1) and minimum (0) intensities within the nucleus in the image. Scale bars represent 10 μm.

**Figure S2.**
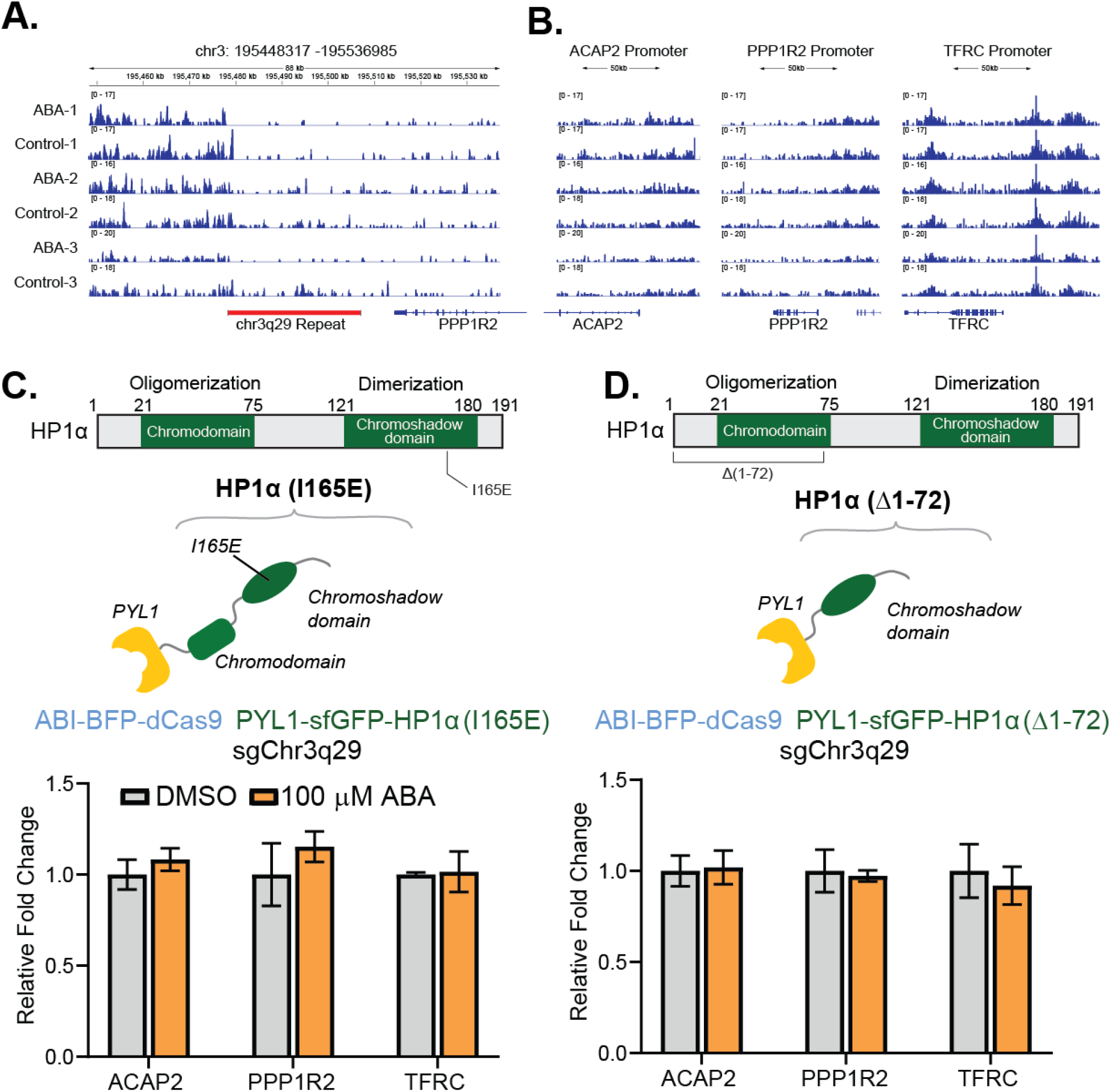
Heterochromatic Properties of CRISPR-EChO at Chr3q29. **(A-B)** H3K9me3 ChIP-seq IGV read coverage histogram tracks of **(A)** a region of Chr3q29 spanning 30 kb on each side of the sgRNA targeted repeat site or **(B)** the promoters of three adjacent genes, mapped in **Fig. 2B**, for three replicate samples of CRISPR-EChO U2OS cells targeting Chr3q29 and treated with 100 μM ABA or DMSO control for two days. Track heights are normalized to total mapped reads for each sample. **(C-D)** RT-qPCR quantifying mRNA levels of the three indicated genes in U2OS cells expressing sgChr3q29 and a mutant HP1α CRISPR-EChO system with either **(C)** a point mutation to abolish dimerization, I165E, or **(D)** deletion of the N-terminus and chromodomains, Δ1-72 a.a. Schematic for mutant effectors shown on top, with fused fluorescent marker not depicted. Cells were treated with DMSO or 100 μM ABA for 5 days before RNA extraction. Data represents mean ± SD for four biological replicates from two independent experiments.

**Figure S3.**
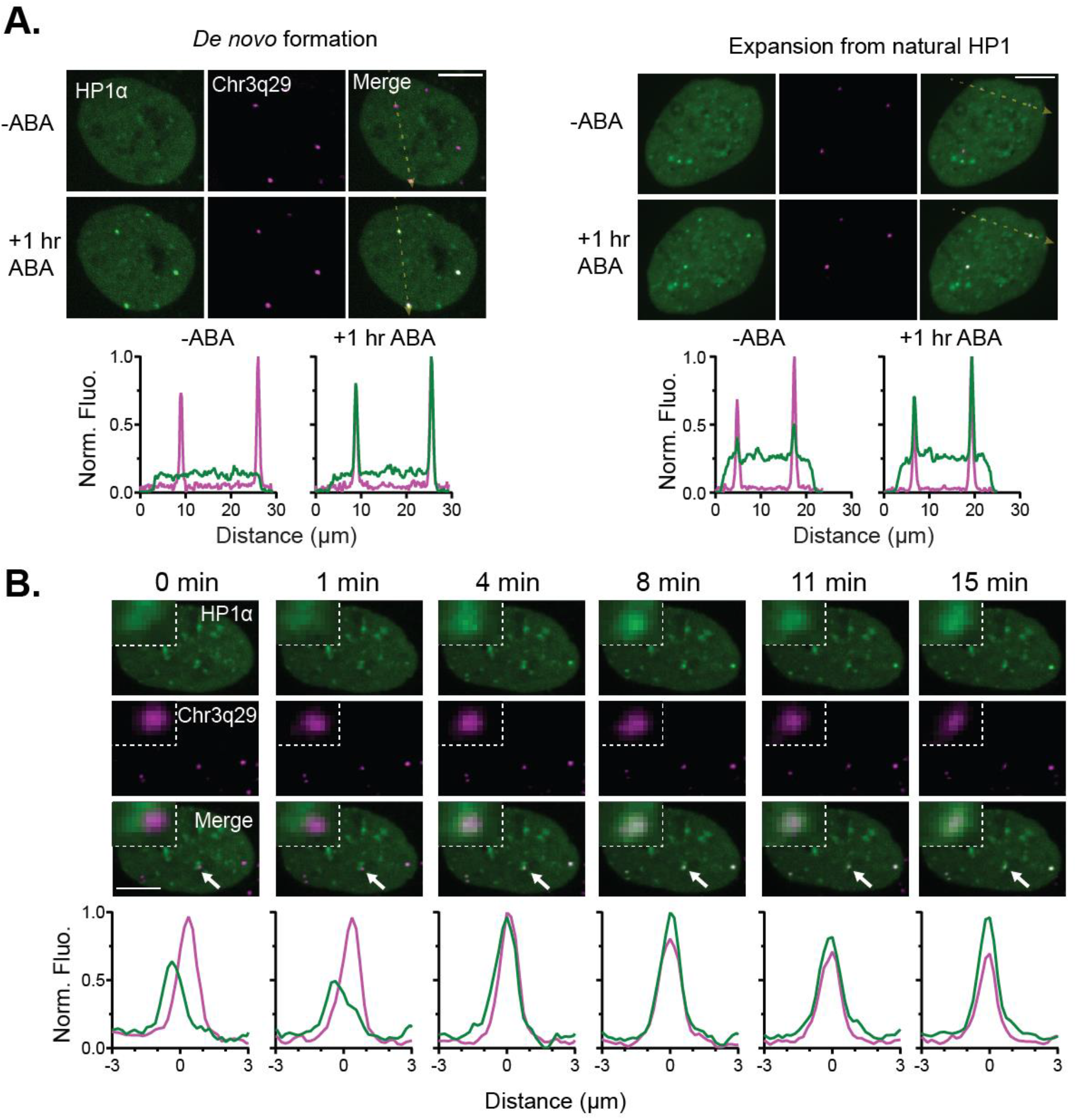
HP1α Assembly and Long-Range Interactions at Chr3q29. **(A)** Representative time-lapse confocal microscopy images of a U2OS cell expressing the HP1α CRISPR-EChO system, MCP-mCherry, and the sgChr3q29-2xMS sgRNA before and after 1 hour 100 μM ABA treatment. Left two panels show a cell that forms HP1α puncta *de novo* at Chr3q29 after ABA addition. Right two panels show a cell with pre-existing HP1α puncta at Chr3q29 that expand after ABA addition. Bottom plots show line scans of fluorescence intensities for each color channel along the yellow dotted line in each Merge image. **(B)** A second representative time-lapse confocal microscopy image (along with **Fig. 3D**) showing incorporation of Chr3q29 into natural HP1α aggregates in the nucleus mediated by CRISPR-EChO. Time 0 min represents image taken immediately before 100 μM ABA addition, and 1 min represents the first image taken of the same cell after addition. Inset image represents 6x magnification of the region indicated by the white arrow. Bottom plots show line scans of fluorescence intensities for each color channel for region indicated by the white arrow. In **(A)** and **(B)**, fluorescence intensities are normalized to the maximum (1) and minimum (0) intensities observed across line scans for all displayed time points. Scale bars represent 10 μm.

**Figure S4.**
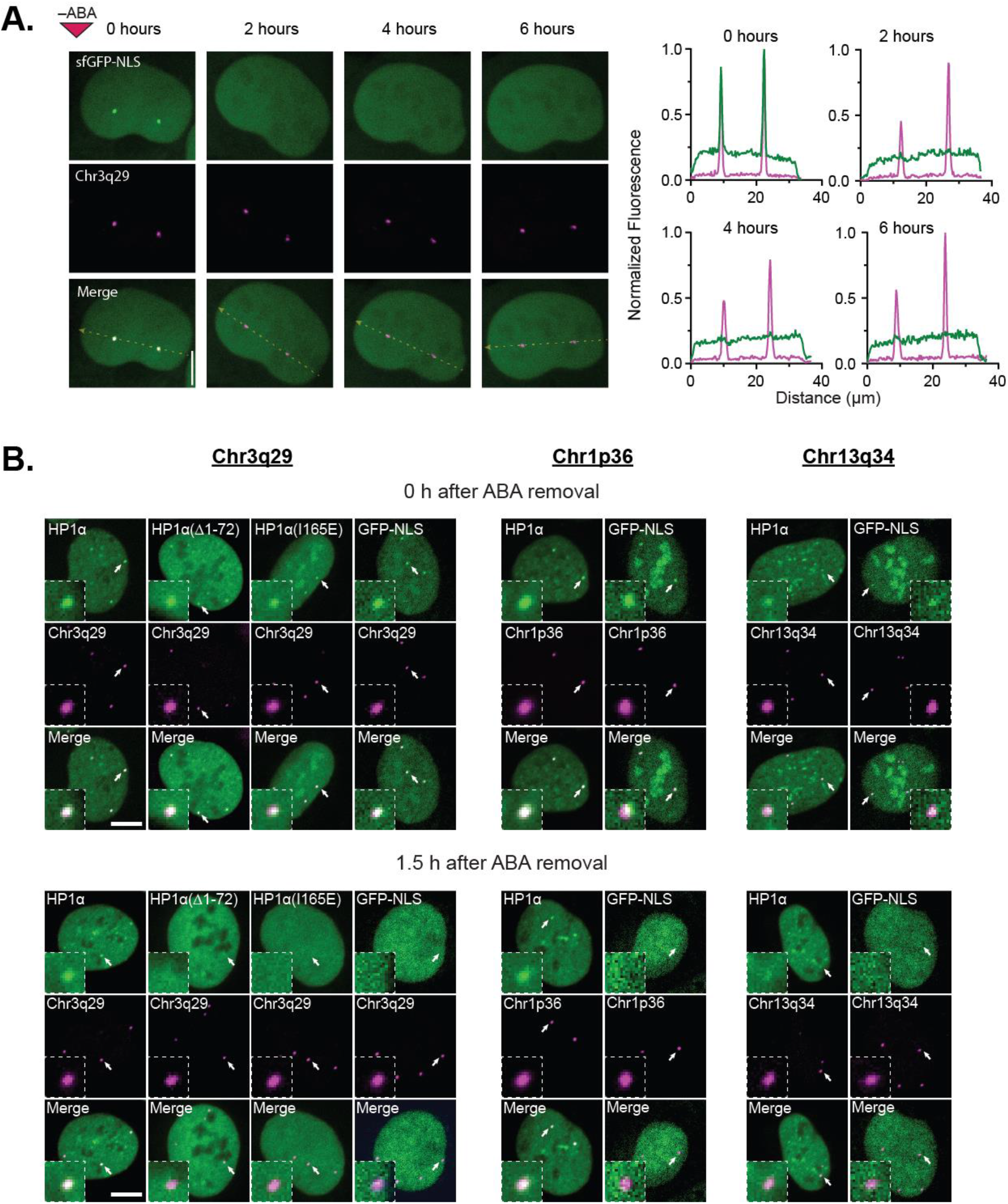
CRISPR-EChO Disassembly Kinetics Upon ABA Removal. **(A)** Representative time-lapse confocal microscopy images of a U2OS cell expressing the CRISPR-EChO system with a PYL1-sfGFP-NLS control effector. Cells were treated for 24 hrs with 100 μM ABA prior to imaging. The 0 hour image represents the first image taken immediately after ABA removal. Bottom plots show line scans of fluorescence intensities for each color channel along the yellow dotted lines in each Merge image. Fluorescence intensities are normalized to the maximum (1) and minimum (0) intensities observed across line scans for all displayed time points. **(B)** Representative confocal microscopy images of U2OS cells expressing the CRISPR-EChO system with various effectors and targeted to Chr3q29, Chr1p36, or Chr13q34. Images were taken immediately after ABA removal or at 1.5 hours after ABA removal. Scale bars represent 10 μm.

**Figure S5.**
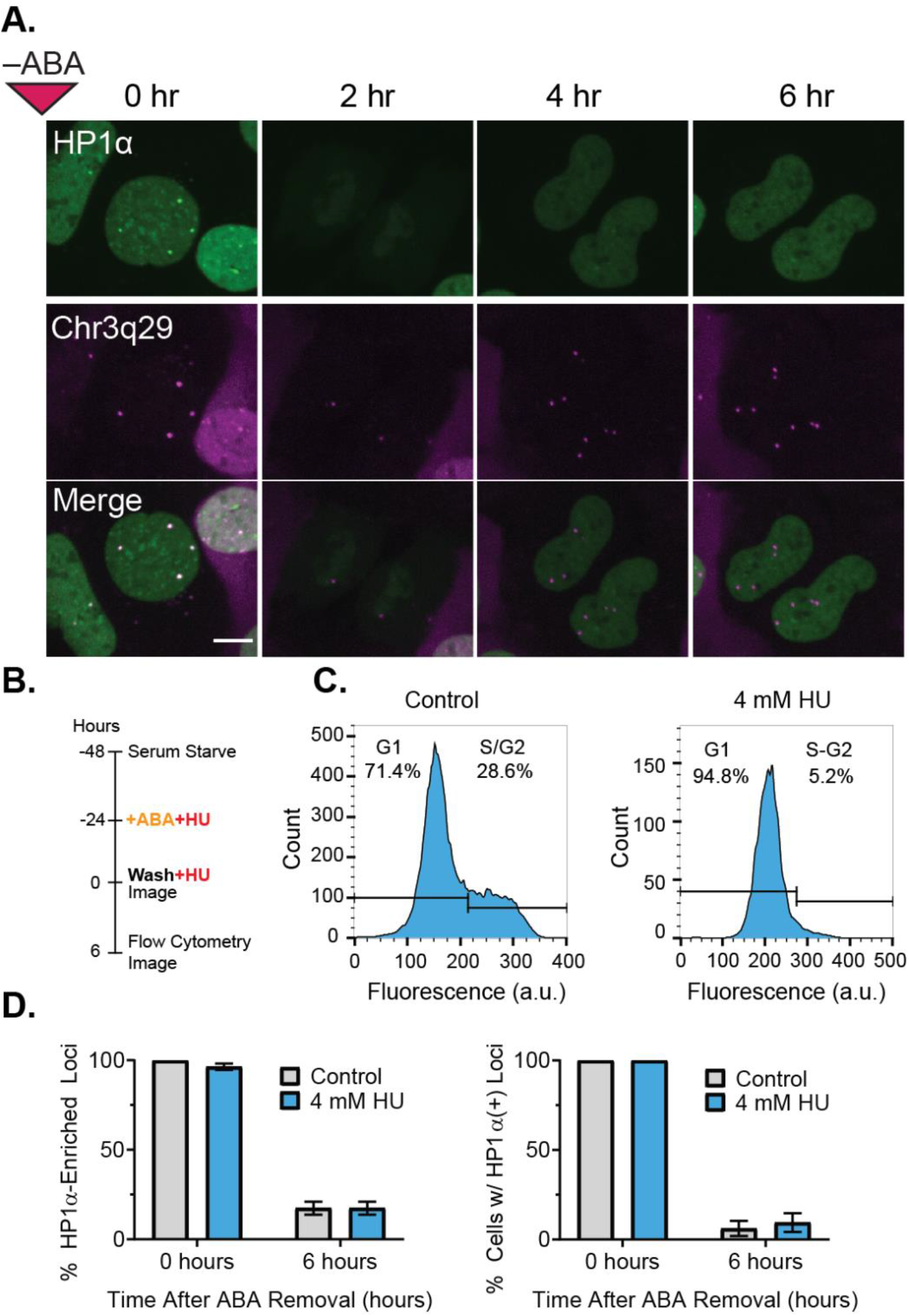
Cell Cycle Arrest Does Not Delay Disassembly of HP1α at Chr3q29. **(A)** Time lapse microscopy images showing a U2OS cell expressing HP1α CRISPR-EChO undergoing mitosis. Bright HP1α puncta observed at Chr3q29 loci at 0 hr are lost during and following mitosis. Cells were treated for 24 hrs with 100 μM ABA prior to imaging. The 0 hour image represents the first image taken immediately after ABA removal. Scale bar represents 10 μm. **(B)** Experimental set-up for serum starvation and hydroxyurea induced cell cycle arrest. Control cells are serum starved but grown in complete media upon ABA addition. **(C)** Flow cytometry quantification of DNA content in Hoechst 33342-stained U2OS HP1α CRISPR-EChO cells with or without hydroxyurea-induced cell cycle arrest. Control cells show 28.6% of cells in S/G2 phase, while 4mM HU-treated cells show 5.2% of cells in S/G2. **(D)** Comparison of HP1α disassembly rates at Chr3q29 between control and cell cycle arrested U2OS HP1α CRISPR-EChO cells. Left panel displays the percentage of all Chr3q29 loci with HP1α-enrichment at the indicated time points (n=109-116 loci). Right panel shows percentage of cells with >50% Chr3q29 loci with GFP enrichment (n=31-34 cells). Error bars represent SEM calculated from Bernoulli distributions.

**Figure S6.**
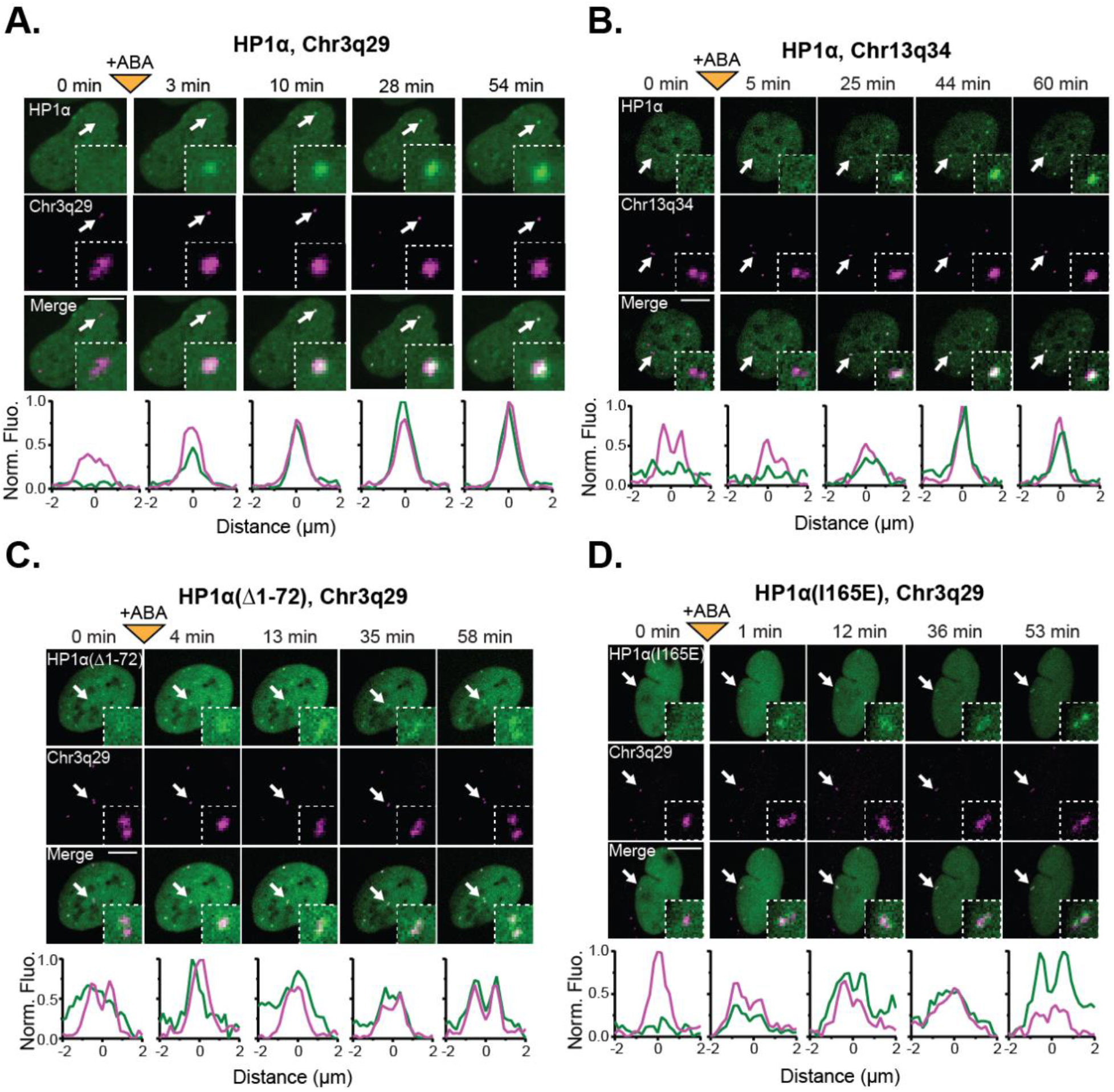
HP1α Recruitment to Tandem Repeat Sites Induces Chromatin Compaction. **(A)** A second representative image (along with **Fig. 5B**) showing compaction of a Chr3q29 locus following 100 μM ABA addition to recruit HP1α. Compaction is concomitant with the assembly of the HP1α puncta at Chr3q29. **(B)** A representative image showing compaction of a Chr13q34 locus following 100 μM ABA addition to recruit HP1α. **(C-D)** Representative images show Chr3q29 loci remaining diffuse following 100 μM ABA addition to recruit mutant **(C)** HP1α(Δ1-72) or **(D)** HP1α(I165E). Inset image represents 6x magnification in **(A)** and 4x magnification in **(B-D)** of the region indicated by the white arrow. For all panels, bottom plots show fluorescence intensities of each color channel for region indicated by the white arrow. Fluorescence intensities are normalized to the maximum (1) and minimum (0) intensities observed across line scans for all displayed time points. Scale bars represent 10 μm.

